# Deep Learning Based Registration of Serial Whole-slide Histopathology Images in Different Stains

**DOI:** 10.1101/2022.05.31.494254

**Authors:** Mousumi Roy, Fusheng Wang, George Teodoro, Shristi Bhattarai, Mahak Bhargava, T Subbanna Rekha, Ritu Aneja, Jun Kong

**Affiliations:** Department of Computer Science, Stony Brook University, NY, 11794, USA; Department of Biomedical Informatics, Stony Brook University, NY, 11794, USA; Department of Computer Science, Federal University of Minas Gerais, Belo Horizonte, 31270-901, Brazil; Department of Biology, Georgia State University, Atlanta, GA, 30303, USA; Department of Clinical and Diagnostic Sciences, School of Health Profession, University of Alabama at Birmingham, Birmingham, AL, 35233, USA; Department of Pathology, JSS Medical College, JSS Academy of Higher Education and Research, Mysuru, Karnataka, India - 570009; Department of Mathematics and Statistics and Computer Science, Georgia State University, Atlanta, GA, 30303, USA; Department of Computer Science and Winship Cancer Institute, Emory University, Atlanta, GA, 30322, USA

**Keywords:** Image registration, Whole-slide image, Multiple stains, Image translation, Deformable vector field

## Abstract

For routine pathology diagnosis and imaging based biomedical research, Whole-Slide Image (WSI) analyses have been largely limited to a 2D tissue image space. For a more definitive tissue representation to support fine-resolution spatial and integrative analyses, it is critical to extend such tissue based investigations to a 3D tissue space with spatially aligned serial tissue WSIs in different stains, such as Hematoxylin and Eosin (H&E) and Immunohistochemistry (IHC) biomarkers. However, such WSI registration is technically challenged by the overwhelming image scale, the complex histology structure change, and the significant difference in tissue appearances in different stains. We propose a novel translation based deep learning registration network CycGANRegNet that spatially aligns serial WSIs stained in H&E and by IHC biomarkers without prior deformation information for the model training. First, synthetic IHC images are produced from H&E slides through a robust image synthesis algorithm. Next, the synthetic and the real IHC images are registered through a Fully Convolutional Network with multi-scaled deformable vector fields and a joint loss optimization. We perform the registration at the full image resolution, retaining the tissue details in the results. Evaluated with a dataset of 76 breast cancer patients with one H&E and two IHC serial WSIs for each patient, CycGANRegNet outperforms multiple state-of-the-art deep learning based and conventional pathology image registration methods. Our results suggest that CycGAN-RegNet can produce promising registration results with serial WSIs in different stains, enabling integrative 3D tissue-based biomedical investigations.

## 1. INTRODUCTION

Histopathology Whole-Slide Images (WSIs) of tissue sections provide high resolution tissue details critical for disease diagnosis and study. Such high resolution WSIs have been largely analyzed in a two-dimension (2D) tissue imaging space by far. As each 2D WSI can only capture information from the tissue cutting plane, it is inevitably subject to information loss and sampling bias problems. Therefore, it is critical to extend such analyses to a three-dimension (3D) tissue space, especially for those studies requiring a definitive tissue characterization. In the meanwhile, such 3D tissue based methods are urged by the rapidly increasing analysis and clinical demand on spatial integration of multi-stained serial WSIs. In particular, there is a strong demand to integrate Hematoxylin and Eosin (H&E) WSIs capturing histology tissue phenotype information with serial Immunohistochemistry (IHC) WSIs visualizing the underlying disease molecular underpinnings for a comprehensive understanding of tumor micro-environment. Apparently, an accurate registration of such serial tissue WSIs is a prerequisite to support such 3D tissue-based analyses. However, WSI registration is technically challenging. Existing methods only achieve a limited success due to the overwhelmingly large image scale, complex histology structure change across adjacent slides, and significant tissue appearance difference in different staining [1].

Recently, the ANHIR challenge was organized to systematically compare the performances of image registration algorithms for microscopy histology images [1]. Most methods described in the ANHIR challenge are based on a rigid registration followed by a non-rigid registration. Rigid registration is determined by RANSAC from feature points whereas non-rigid registration is performed by local affine transformation, demons algorithms, or interpolations. For example, a two-stage image registration method is proposed where the registration is formulated as an optimization problem with an objective function defined by Normalized Gradient Fields (NGF) based distance measure [2]. In another study, a fine tuning method is proposed based on integrated landmark evaluation by texture and spatial proximity measures [3]. In a study on multi-stained histology image registration, a method is developed by the feature-based affine registration, rotation alignment followed by nonrigid alignment with Demons algorithm [4]. Moreover, a modified SIFT method is applied to image registration with color interpolation [5]. WSIReg is another method for WSI registration based on elastix [6]. However, all these methods require feature matching that highly depends on feature engineering and parameter tuning.

With the emergence of deep learning methods, it is possible to leverage the image-to-image translation for spatial alignment of images in different appearances [7, 8, 9]. Translation-based approaches use Generative Adversarial Network (GAN) to translate images from one modality (e.g. H&E) to another (e.g. IHC), simplifying the image registration task. Although much simplified, such an analysis still presents two significant challenges. First, H&E and IHC WSIs are unpaired data as each pathology tissue slide is stained only once in most clinical practice. Second, it is time-consuming and financially costly to have accurate landmark pair annotations from serial WSIs for registration. Recently, CycleGAN with the cycle consistent loss has been developed to learn an image-to-image mapping between two domains from unpaired data [10]. The cycle consistency loss for the adversarial training process forces the generator to find an accurate mapping between two different domains with unpaired data. By this approach, synthetic slice-wise computed tomography data has been produced from magnetic resonance images [11]. In another study on thoracic and abdominal organs, a mono-modal image registration with CycleGAN presents a comparable performance to the multimodal deformable registration with paired image data [7]. Furthermore, a deformation field is used for MRI-CT registration in a dual-stream fashion with CycleGAN [12]. In this work, the MIND based loss term added to CycleGAN loss describes the local image structure. As such a loss term is computed with gray-scale images, and such a method cannot be applied to multi-stained pathology image data directly. In another study, a CycleGAN based image generation method generates IHC histology images from H&E images without any annotation [13] where the class-related information is added as an additional input patch channel. Therefore, this method is not directly applicable to our study with multistained pathology image data.

In this paper, we present a new translation based deep learning registration approach (CycGANRegNet) for serial WSIs in different stains. It consists of an image translation and an image registration module. The image translation module produces synthetic IHC slides (i.e. synIHC images) from real H&E slides through a robust image synthesis algorithm. With Fully Convolutional Network (FCN) model [14] as a building block, the synIHC and the real IHC image pairs are registered through a multi-scale based FCN registration model. The major contributions of this paper are summarized in multiple folds:

- We develop a modified CycleGAN method to generate synthetic IHC pathology images (i.e. synIHC) from unpaired serial H&E pathology slides. To enhance the image stain translation ability, we propose to adopt a perceptual loss in the CycleGAN loss function, resulting in a better image mapping from H&E to synthetic IHC images. Such an image translation enables a better registration between synIHC and real IHC images.
- We extend the original FCN model to a multi-scaled architecture for registration. Our proposed multi-scale FCN uses a coarse-to-fine multi-scale deformable image registration strategy that combines the Displacement Vector Fields (DVFs) at multiple resolutions for better image alignment.
- CycGANRegNet is an unsupervised registration approach and can be efficiently trained without ground truth image deformation information.
- Instead of resizing images to a lower resolution [1], we recover the WSI registration results with patch-based image registration results at the highest image resolution. Therefore, high resolution tissue details from WSIs are retained.

## 2. METHODS

We develop a deep learning based model CycGANRegNet to register serial IHC to H&E histopathology images for molecular biomarker and pathology hallmark integration. It is an end-to-end deep learning process in two stages. First, we develop an image translation module with a modified Cycle-GAN to translate real H&E references to synthetic IHC image patches (i.e. synIHC). Next, we develop a multi-scale FCN in the image registration module to estimate the spatial mapping from the real IHC to the synthetically produced synIHC image patches. Then the real IHC image is transformed to the reference H&E image space via Spatial Transformation Network (STN) [15]. Finally, individual registered image patches are spatially assembled to recover registered WSI blocks. We present the overall schema of CycGANRegNet in Figure 1.

**Fig. 1.**
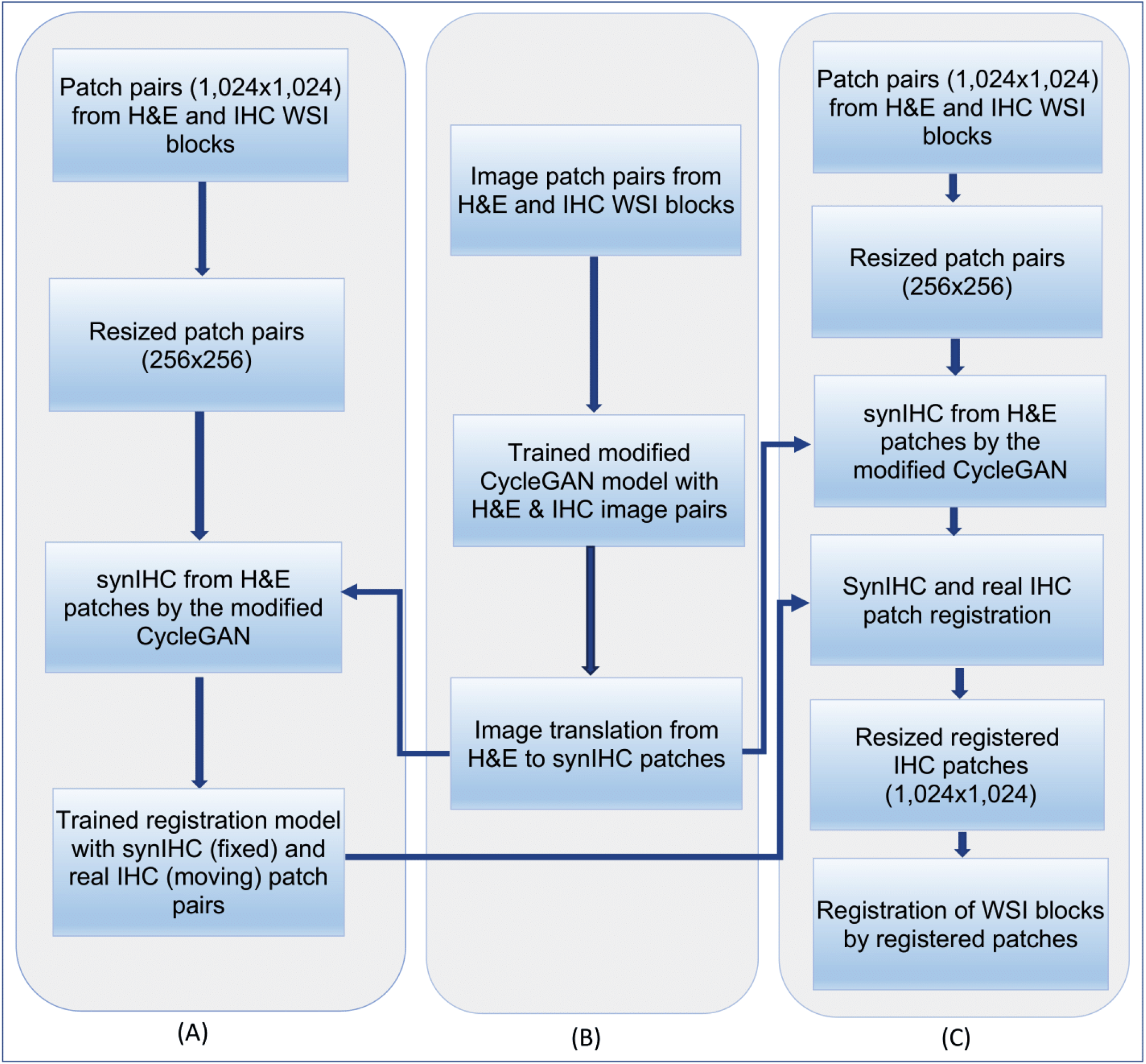
The overall schema of the CycGANRegNet. (A) Patch-based H&E-IHC image registration; (B) Translation from H&E to synIHC images; and (C) WSI block registration.

### 2.1. Unpaired Image Translation

Although serial slides in different stains look similar at the global tissue level (Supplemental Figure 1), they are unpaired at the pixel level. Although CycleGAN can be applied to unpaired data [13, 10, 16], we propose a modified CycleGAN for an enhanced H&E-IHC image translation. Illustrated in Figure 2, the modified CycleGAN consists of an encoder and a decoder. Both have the same network structure that includes a generator and a discriminator. The generator translates an image between stain domains and the discriminator assesses the generated image quality. The modified CycleGAN model consists of two generators *G_HE_* and *G_IHC_*. The generator *G_HE_* translates a real IHC to a synthetic H&E image (i.e. synHE), while *G_IHC_* translates a synHE to a synthetic IHC image (i.e. synIHC) denoted by the red arrows. Similarly, the reverse translation goes from H&E to synIHC and then from synIHC to synHE (i.e. the black arrows). Each generator module consists of two-dimensional fully convolutional networks with nine residual blocks and two fractionally strided convolution layers [19, 17]. Additionally, the model has two discriminators *D_HE_* and *D_IHC_* for distinguishing between translated (i.e. synHE) and real H&E images, and between translated (i.e. synIHC) and real IHC images, respectively. Each discriminator has a fully convolutional architecture to predict if overlapping image patches of size 70 × 70 by pixels are real or synthetic [18]. The leaky ReLU activation function with a factor 0.2 is used. All data are normalized with the instance normalization.

**Fig. 2.**
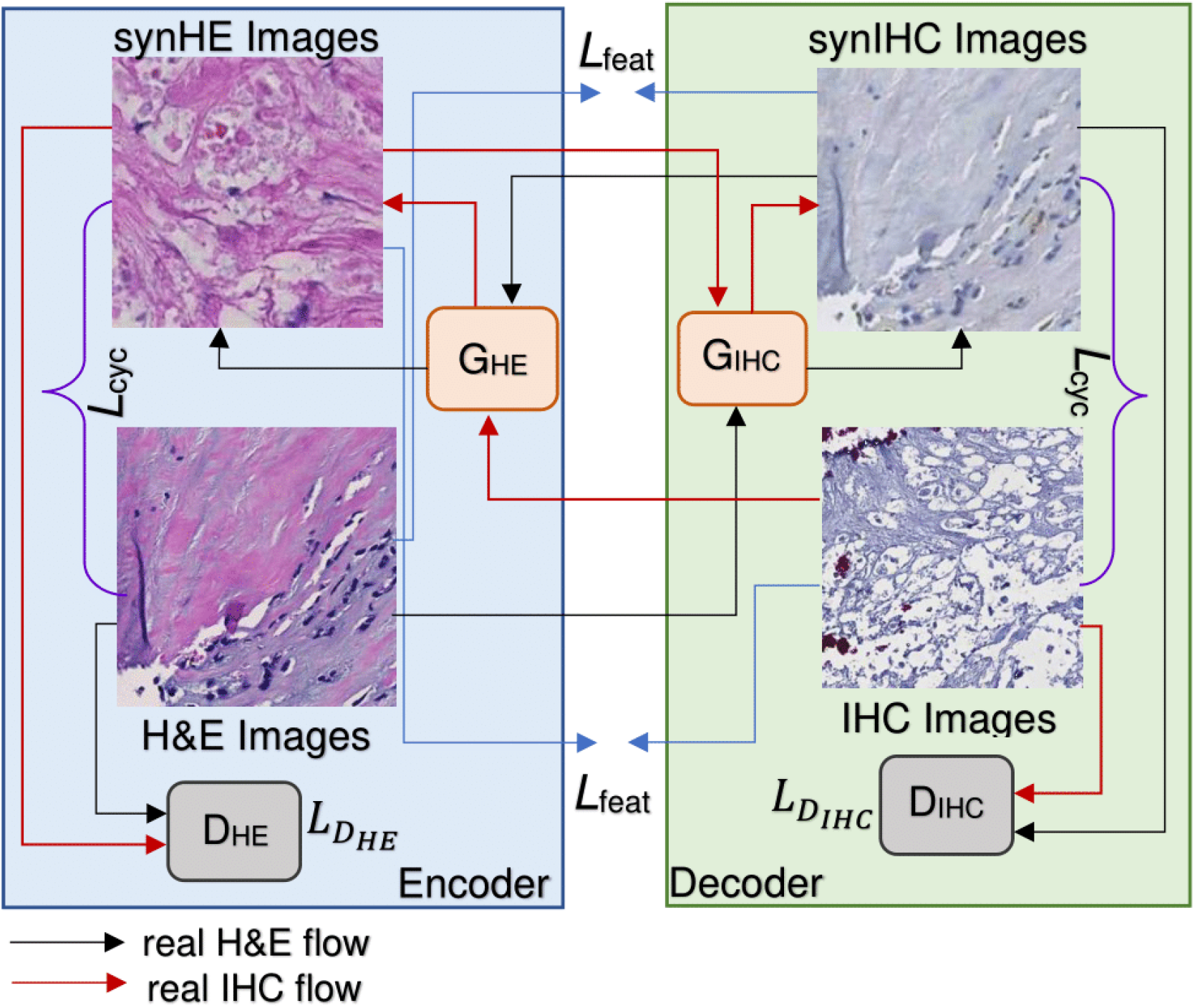
The overall schema of the modified CycleGAN with the forward and backward translation information flow.

The CycleGAN training loss includes the adversarial loss (i.e. *L_D_HE__* and *L_DI_HC__*) from two discriminators and the cycle-consistent loss (*L_cyc_*) [10]. Although the cycle-consistent loss is designed to prohibit the generators from generating images not related to the inputs, this loss by itself is not sufficient to enforce either feature or structural similarity between translated and real images. To address this problem, we adopt the VGG-16 based perceptual loss function as an additional constraint in the CycleGAN loss function. Such a perceptual loss addition helps regularize the tissue content and the stain style discrepancies, as it can measure high-level perceptual and semantic differences between each image pair [19]. Such a perceptual loss is produced by a deep convolutional neural network denoted as *ϕ*, specifically a 16-layer VGG network [20] pre-trained on the ImageNet dataset [21]. The network *ϕ* forces the transformed network output image 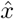 to have similar feature representations as the input real image *x*. The feature reconstruction loss 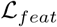 can be written as:

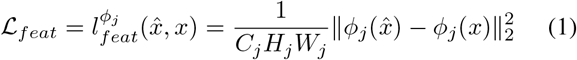

where *ϕ_j_*(*x*) is the activation map of size *C_j_* × *H_j_* × *W_j_* at the *j*-th convolutional layer of network *ϕ* when processing image *x*. Although the image transformation network trained by the feature reconstruction loss encourages the output image 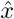 to be perceptually similar to the target image *x*, it does not force them to match each other exactly. The total loss 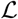 of our modified CycleGAN can be written as:

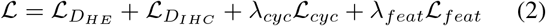

where 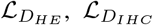 and 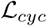 are defined in prior studies [10]. *λ_cyc_* and *λ_feat_* are weights for loss term 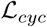 and 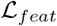, respectively.

### 2.2. Multi-scaled Image Patch Registration

Numerous state-of-the-art histopathology image registration methods are either based on SIFT-feature matching or MIND based Demons algorithms. Some learning based methods (e.g. TUB from ANHIR challenge [1]) have been proposed with limited performance. In this study, we leverage FCN based deep learning model for histopathology image registration due to its known promising performance [14]. As multiple prior studies on flow estimation have shown the effectiveness of multi-scale strategy [22, 23], we use the FCN and the multi-scale strategy as building blocks for the proposed registration method. Specifically, we create a multiscale FCN model with multi-scale Deformable Vector Fields (DVFs) that enable a coarse-to-fine multi-scale deformable image registration. For a better deformation estimation, three DVF estimation models at three different image resolutions are used to define the multi-scale spatial mappings. Our registration pipeline is demonstrated in Figure 3.

**Fig. 3.**
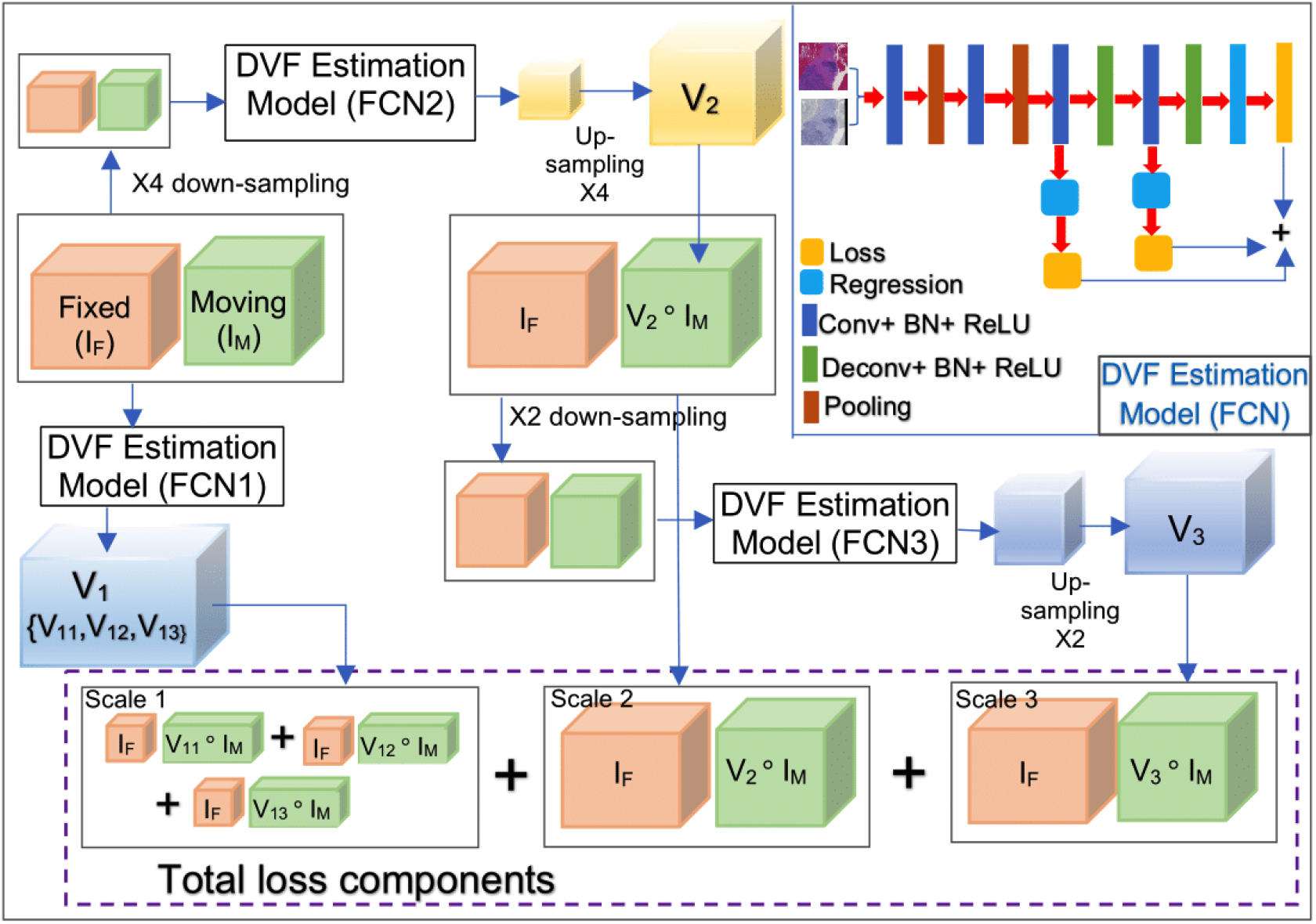
The overall architecture of the developed multi-scale FCN model is presented with detailed illustrations of FCN layers for the multi-scale DVF estimation and multiple loss function components in the total loss.

For image registration, a moving *I_M_* and a fixed *I_F_* image are provided to our multi-scale FCN model as inputs. For model training, a pair of real moving and fixed images are concatenated and provided to the first DVF estimation model to estimate the DVF *V*_1_ at scale-1. *V*_1_ has three different components (i.e. *V*_11_, *V*_12_ and *V*_13_) that are produced by three different layers of the FCN model. The resulting warped moving images are *V*_11_ ◦ *I_M_*, *V*_12_ ◦ *I_M_*, and *V*_13_ ◦ *I_M_*, respectively, where ◦ is the operator applying a DVF to an image. Next, the fixed I_*F*_ and moving *I_M_* image pairs are concatenated and down-sampled by a factor of four. The resulting image pairs are provided to the second DVF estimation model for DVF estimation at scale-2. The resulting DVF is up-sampled to match the original input image size and denoted as *V*_2_. Similarly, *V*_2_ is applied to the input moving image *I_M_* to generate the warped moving image *V*_2_ ◦ *I_M_*. In the next step, the warped moving image *V*_2_ ◦ *I_M_* at scale-2 and the input fixed image *I_F_* are concatenated and down-sampled by a factor of two. The resulting image pairs are provided to the third DVF estimation model for the residual DVF estimation. The resulting DVF is up-sampled to the original input image size and denoted as *V*_3_ at scale-3. Finally, *V*_3_ is used to deform the moving image *I_M_* for the warped image *V*_3_ ◦ *I_M_*. We train all three DVF estimation models simultaneously to minimize the joint loss at multiple scale levels, achieving an optimal end-to-end performance. Our formulated total loss function can be represented as follows:

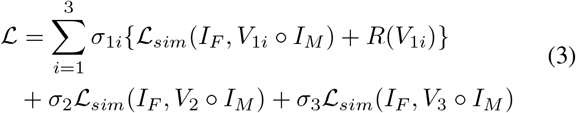

where 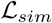 is the similarity loss measured by the Negative Normalized Cross-Correlation (NCC) [24], penalizing the differences in appearance between the fixed and moving images. Note *σ*_11_, *σ*_12_ and *σ*_13_ are weights of the similarity loss metrics at scale level 1, while *σ_2_* and *σ*_3_ are weights at scale levels 2 and 3, respectively. *R*(*V*) is a total variation based regularizer that makes the transformation spatially smooth and physically plausible [25].

After the weight initialization, all weights are updated by the joint training of three DVF estimation models in an end-to-end manner for the harmonic minimization of the composite loss. With displacement vectors between the fixed and moving image pairs, we use Spatial Transformer Network (STN) [15] to deform the moving image [14]. To make the resulting registered images retain more tissue details, we utilize the Enhanced SRGAN (ESRGAN) model [26] in the postprocessing step. Figure 4 demonstrates the registered images by our proposed multi-scale FCN model with and without the post-processing step. There is a noticeable difference in registered images with and without such post-processing. Additionally, such post-processing step also improves the registration performance as suggested by all performance metrics in Table 2.

**Fig. 4.**
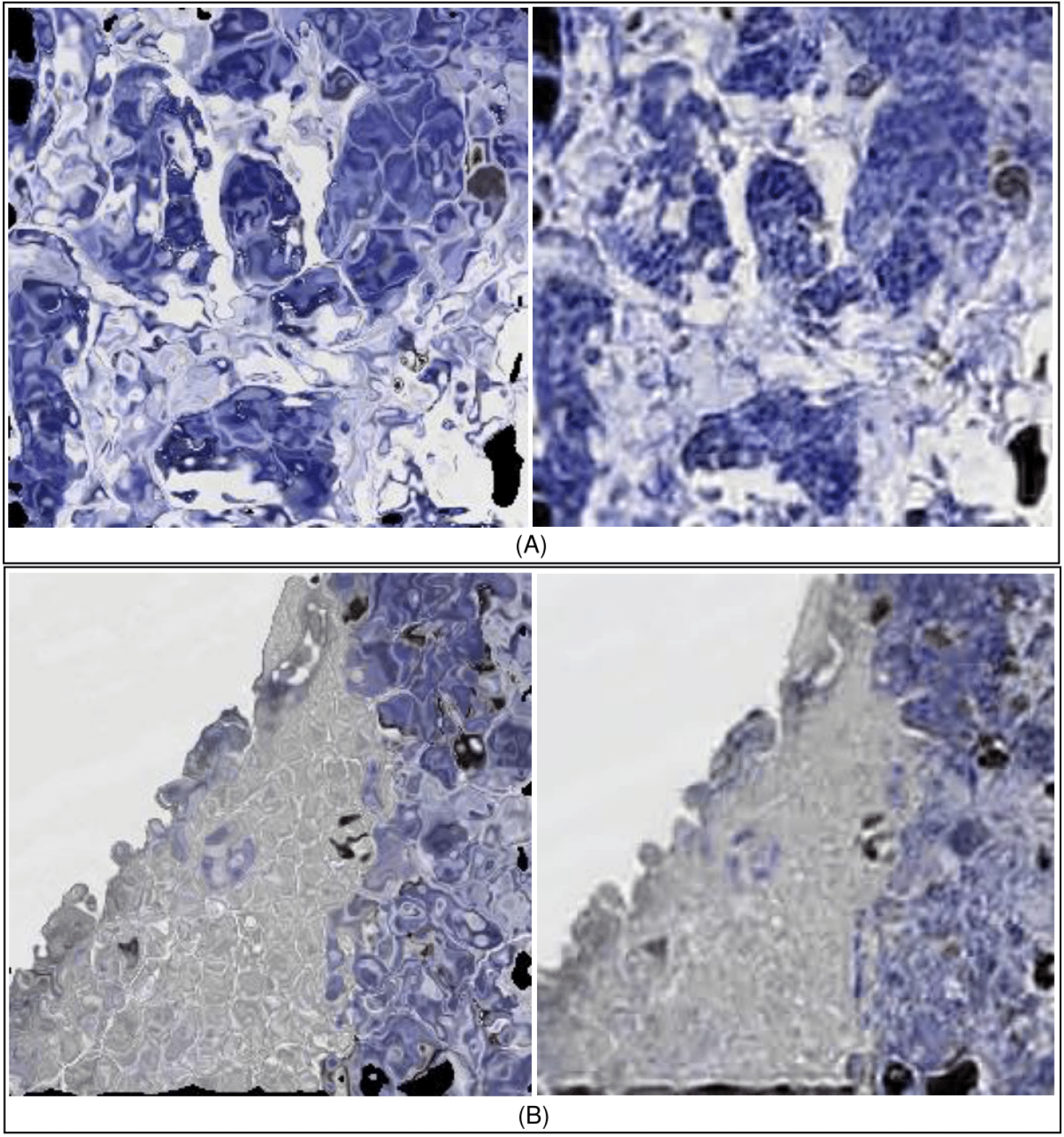
Registration results from our multi-scale FCN model with and without the ESRGAN based post-processing. Two registered image patches from (A) our testing, and (B) validation data (Left) without and (Right) with the post-processing are illustrated.

**Table 1.** Quantitative comparison of image translation performance between the original and our modified CycleGAN model on the testing and validation data.

| Testing Data |  |  |  | Validation Data |  |  |  |
| --- | --- | --- | --- | --- | --- | --- | --- |
|  | RMSE | SSIM | PSNR |  | RMSE | SSIM | PSNR |
| CycleGAN<br>(H&E → Ki67) | 15.200 | 0.934 | 29.928 | CycleGAN<br>(H&E → PHH3) | 17.641 | 0.905 | 28.528 |
| Our<br>(H&E → Ki67) | **14.414** | **0.935** | **30.387** | Our<br>(H&E → PHH3) | **17.210** | **0.911** | **28.754** |
| CycleGAN<br>(H&E ← Ki67) | 13.088 | 0.955 | 31.215 | CycleGAN<br>(H&E ← PHH3) | 15.114 | 0.928 | 29.956 |
| Our<br>(H&E ← Ki67) | **12.123** | **0.959** | **31.867** | Our<br>(H&E ← PHH3) | **14.430** | **0.935** | **30.282** |

**Table 2.** Image patch registration performance on the testing and validation data.

| Method Name | dataset | Testing Data |  |  | Validation Data |  |  |
| --- | --- | --- | --- | --- | --- | --- | --- |
|  |  | NCC | SSIM | NMI | NCC | SSIM | NMI |
| DirNet | real | 0.2045 | 0.2307 | 1.0358 | 0.2030 | 0.2413 | 1.0344 |
|  | syn-1 | 0.2644 | 0.3320 | 1.0356 | 0.2613 | 0.3161 | 1.0334 |
|  | syn-2 | 0.2977 | **0.3505** | 1.0374 | 0.2553 | **0.3443** | 1.0324 |
| VoxelMorph<br>UNet | real | 0.0231 | 0.1922 | 1.0329 | 0.0346 | 0.2078 | 1.0324 |
|  | syn-1 | 0.0755 | 0.3164 | 1.0314 | 0.0910 | 0.2828 | 1.0321 |
|  | syn-2 | 0.1063 | 0.3384 | 1.0347 | 0.1306 | 0.3243 | 1.0260 |
| FCN | real | 0.2165 | 0.2168 | 1.0373 | 0.1742 | 0.2266 | 1.0362 |
|  | syn-1 | 0.2508 | 0.3119 | 1.0363 | **0.2688** | 0.3050 | 1.0339 |
|  | syn-2 | 0.3057 | 0.3356 | **1.0392** | 0.2683 | 0.3356 | 1.0340 |
| Our Multi-scale FCN | real | 0.1971 | 0.2214 | 1.0370 | 0.1776 | 0.2344 | 1.0360 |
|  | syn-1 | 0.2582 | 0.3096 | 1.0370 | 0.2321 | 0.2914 | **1.0362** |
|  | syn-2 | **0.3161** | <u>0.3403</u> | <u>1.0387</u> | <u>0.2625</u> | <u>0.3375</u> | 1.0343 |

### 2.3. WSI Block Registration

Due to the limited GPU memory size, deep learning methods cannot process giga-pixel WSIs at the full histopathology image resolution all at a time. Therefore, each WSI is partitioned into image blocks of size 8, 000 × 8, 000 pixels for tissue pre-alignment. Each H&E block is next translated to a synIHC block by the developed modified CycleGAN. Real IHC and synIHC image blocks are next divided into image patches of size 1, 024 × 1, 024 to retain sufficient tissue information for registration. The resulting synIHC and real IHC patch pairs are further resized to 256 × 256 pixels for the deep learning model training and prediction. After registration, the registered real IHC image patches are resized back to 1, 024 × 1, 024 size and spatially assembled for WSI block registration [27].

## 3. EXPERIMENTAL RESULT

### 3.1. Dataset and Implementation

This study cohort consists of 76 Neoadjuvant Chemotherapy (NAC) treated Triple Negative Breast Cancer (TNBC) patients from Emory Decatur Hospital. Formalin-fixed paraffin-embedded serial biopsy sections from each patient are H&E and immunohistochemically stained with Ki67 as a biomarker for cell proliferation and Phosphohistone H3 (PHH3) for the mitotic activity. In total, we assess our model on 228 WSIs, with three serial WSIs from each patient. After the image pre-alignment by the global affine spatial transformation at a low image resolution, the resulting transformation is mapped to the full image resolution level. The pre-aligned tissue regions at the full image resolution level are next partitioned into 1,023 WSI blocks of size 8, 000 × 8, 000 pixels by each stain. The pre-aligned WSI blocks are further partitioned into non-overlapping image patches of size 1, 024 × 1, 024 pixels and resized to 256 × 256 to make the image size appropriate for deep learning models. Patches containing more than 30%background pixels are excluded from further analyses. This process results in 60,000 image patches that are randomly divided into training, validation and testing cohorts by patients at the ratio of 80:10:10.

Our model is first tested with serial slides in H&E and of Ki67 IHC biomarker, followed by additional validation with serial slides in H&E and of PHH3 IHC biomarker. We compare our model with multiple state-of-the-art methods using both ‘real’ and ‘synthetic’ datasets. The ‘real’ dataset includes pairs of H&E and real IHC images, while the ‘synthetic’ dataset consists of real and synthetic (i.e. synIHC) IHC image pairs with synKi67 for testing and synPHH3 for validation, respectively. Note the dataset with synthetic IHC images by the CycleGAN is labeled ‘syn-1’, whereas that with synthetic IHC images by our modified CycleGAN model is labeled ‘syn-2’. The developed CycGANRegNet is implemented with the open-source deep learning library Tensorflow [28], while experiments are executed on GPUs(i.e. Tesla K80 and V100) with CUDA 9.1. Adam optimization algorithm [29] with learning rate 0.0001 is used to train both image translation and image registration model. The modified CycleGAN is trained for up to 2,00,000 iterations. Loss weights *λ_cyc_* and *λ_feat_* are set to 1. All other parameter settings are suggested by the original CycleGAN work [10]. The values of registration loss weights *σ*_11_, *σ*_12_, *σ*_13_, *σ*_2_ and *σ*_3_ are empirically set to 0.9, 0.6, 0.3, 0.05 and 0.05, respectively.

### 3.2. Evaluation of Image Translation

Both CycleGAN and our modified CycleGAN for image translation are evaluated and compared at the patch and WSI block level. Representative image patches and WSI blocks after translation are demonstrated in Figure 5 and Figure 6, respectively. In addition to the qualitative assessments, we quantitatively evaluate the translated IHC image quality by Root Mean Square Error (RMSE), Structural Similarity Index Measure (SSIM), and Peak Signal-to-Noise Ratio (PSNR) with the maximum value range 40dB [30, 31, 12]. In our analysis pipeline, the trained generator *G_IHC_* and *G_HE_* are used to translate a H&E to a synIHC image, and then back from the synIHC to the synHE (i.e. black arrows in Figure 2). The similarity between the real H&E and resulting synHE image is quantitatively evaluated and presented in Table 1. Both forward (i.e. H&E to synIHC and synIHC to synHE) and reversed (i.e. IHC to synHE and synHE to synIHC) translation performances by the original and our modified CycleGAN model are presented. In our study, the image translation between H&E and Ki67 is for testing, while the H&E-PHH3 translation is for validation purpose. Suggested by experimental results, our proposed modified CycleGAN presents a consistent superior performance to the original CycleGAN by all evaluation metrics in both forward and reversed translation directions. Additionally, the translated synIHC image patches are spatially combined to generate the translated synIHC WSI blocks as illustrated in Figure 6. Both qualitative and quantitative experiments with the testing and validation data suggest an enhanced image translation performance by our modified CycleGAN.

**Fig. 5.**
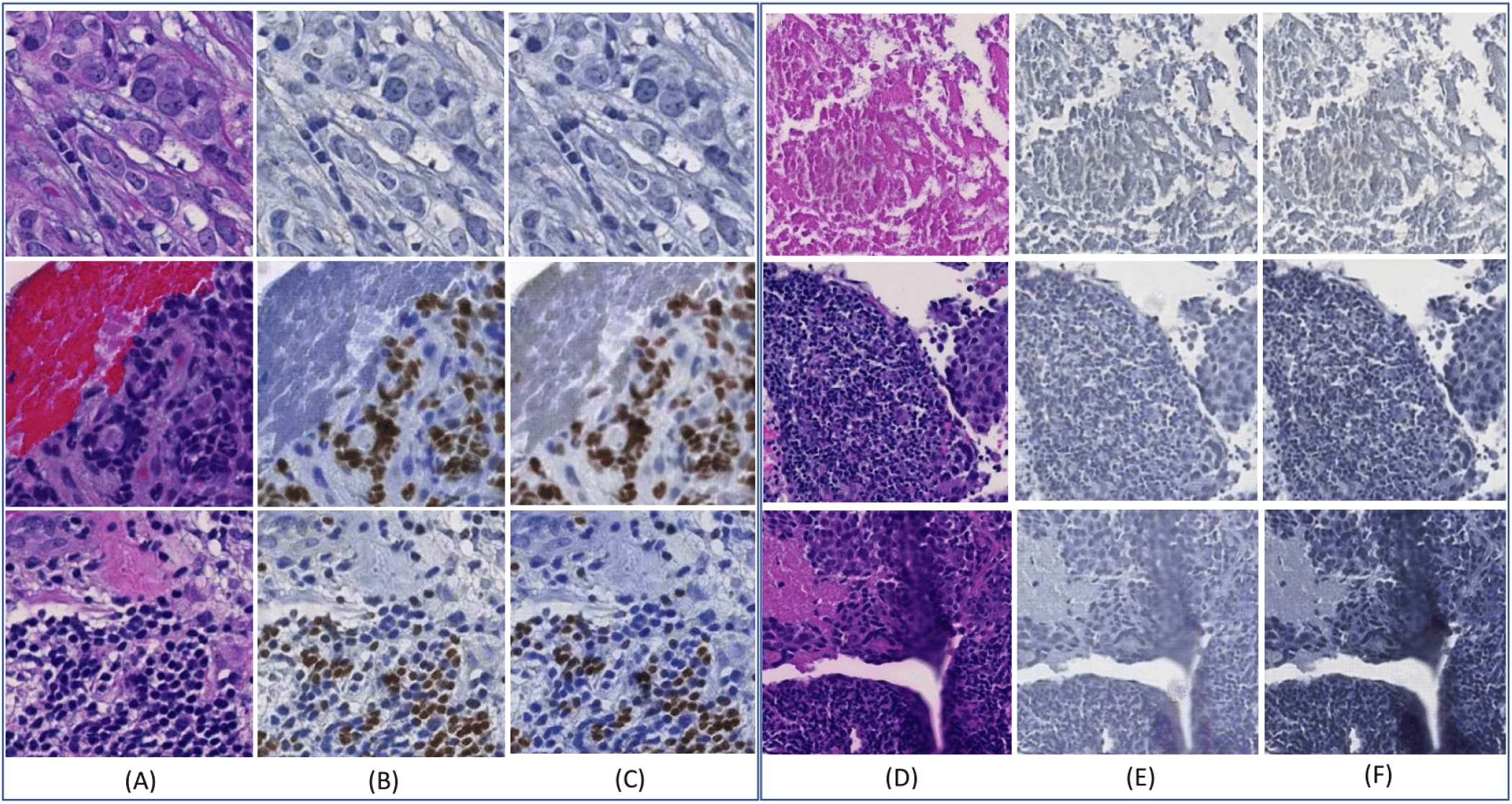
Representative image patch translation results with the testing (left) and validation data (right). (A) Real H&E image patch; (B) synKi67 patch by CycleGAN; (C) synKi67 patch by our modified CycleGAN for the testing data; (D) Real H&E patch; (E) synPHH3 patch by CycleGAN; (F) synPHH3 patch by our modified CycleGAN for the validation data.

**Fig. 6.**
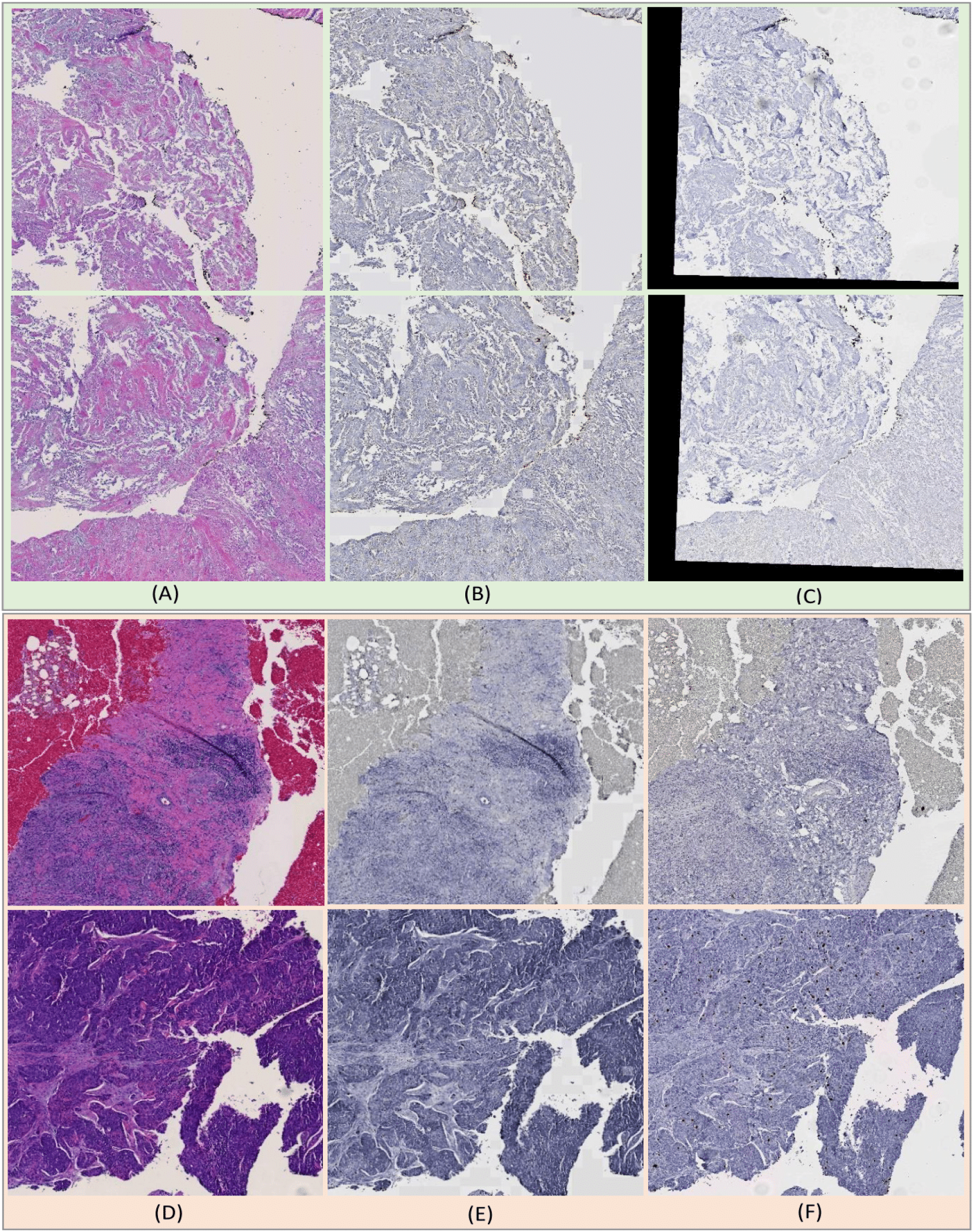
Representative WSI block translation results with the testing (top) and validation data (bottom). (A) Real H&E WSI blocks; (B) synKi67 WSI blocks by our modified CycleGAN; (C) Real Ki67 WSI blocks; (D) Real H&E WSI blocks; (E) synPHH3 WSI blocks by our modified CycleGAN; (F) Real PHH3 WSI blocks.

### 3.3. Evaluation of Image Patch Registration

After the image translation, the resulting synIHC and real IHC WSI blocks are first pre-aligned by a global intensity-guided rigid transformation [32]. Next, CycGANRegNet takes rigidly registered synIHC and real IHC WSI blocks as inputs and fine tunes image alignment by our proposed multi-scale FCN model. To evaluate the registration performance, we apply our method to the ‘real’, ‘syn-1’ and ‘syn-2’ dataset from both the testing data with H&E and Ki67 image pairs and validation data with H&E and PHH3 image pairs. For performance evaluation, our proposed multi-scale FCN model is compared with multiple state-of-the-art registration methods, including deep learning based DirNet [33], FCN [14], VoxelMorph Unet [34] and the conventional image registration method Elastix [35].

As the H&E and IHC WSI blocks are pre-aligned by the global affine transformation before registration methods for comparison in this study, the registration effect on certain image patches is not visually salient. To manifest the method efficacy, we thus manually deform the moving images by affine and elastic transformations, resulting in deformed moving images with significant deformations. Additionally, diverse shear transformations with the rotation angle range of [-40, 30] and elastic deformations are applied to moving images. With such artificial transformation operations, 20, 000 synthetically deformed image patches of size 256 × 256 are derived from 1,023 WSI blocks for the training purpose. Typical registration results by state-of-the-art deep learning registration models with moving images warped by the shear and the elastic deformation are presented in Figure 7 and Figure 8, respectively. By visual assessments, FCN and our proposed multi-scale FCN model present noticeably better registration results than other deep learning models.

**Fig. 7.**
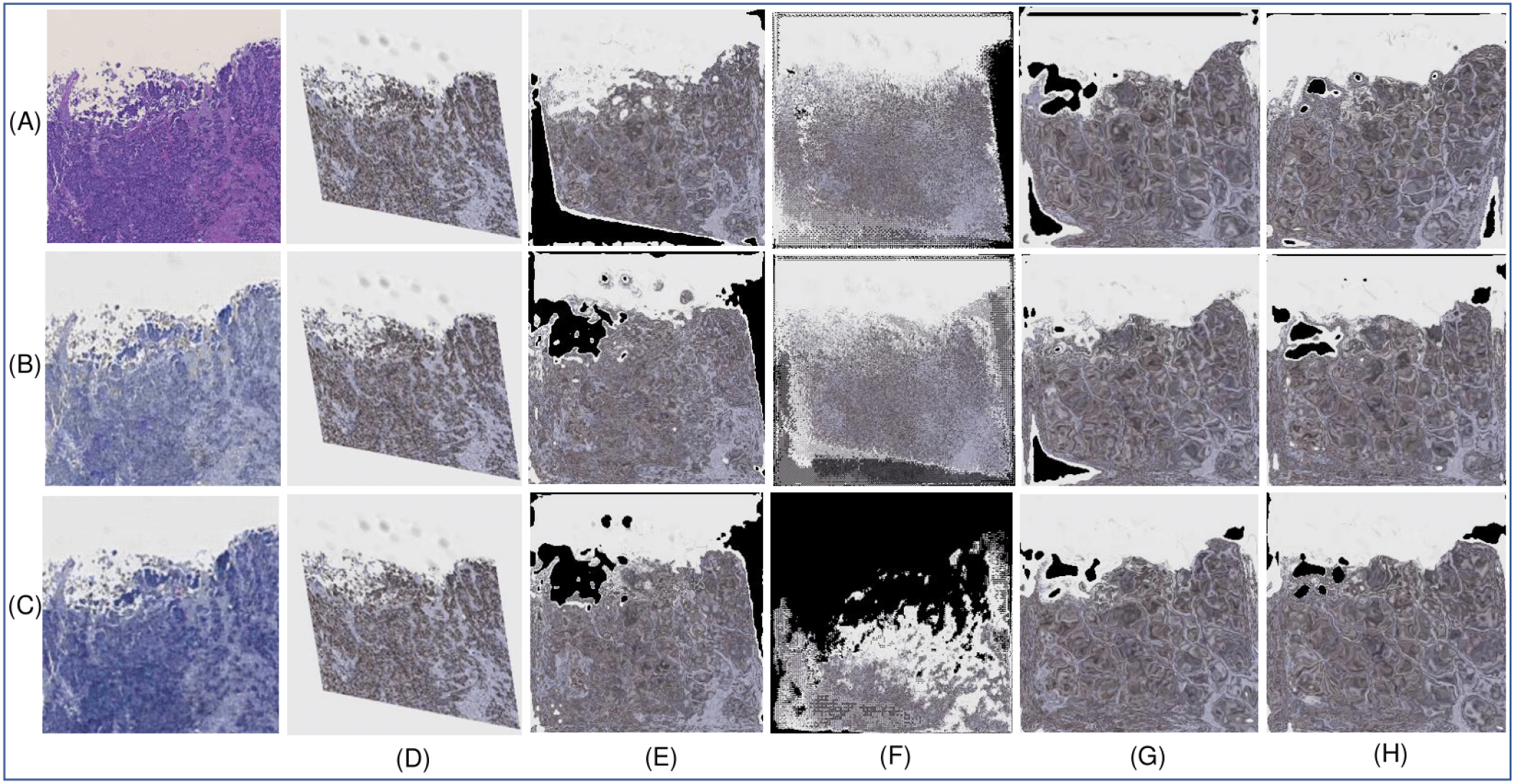
Patch based registration performance with the moving image artificially warped by a shear transformation. (A)Fixed real H&E image; (B)The resulting synIHC image from (A) by CycleGAN; (C)The resulting synIHC image from (A) by our modified CycleGAN; (D) Moving images after manual transformations; Registration results by (E)DirNet, (F)VoxelMorph Unet, (G)FCN, and (H)our proposed multi-scale FCN.

**Fig. 8.**
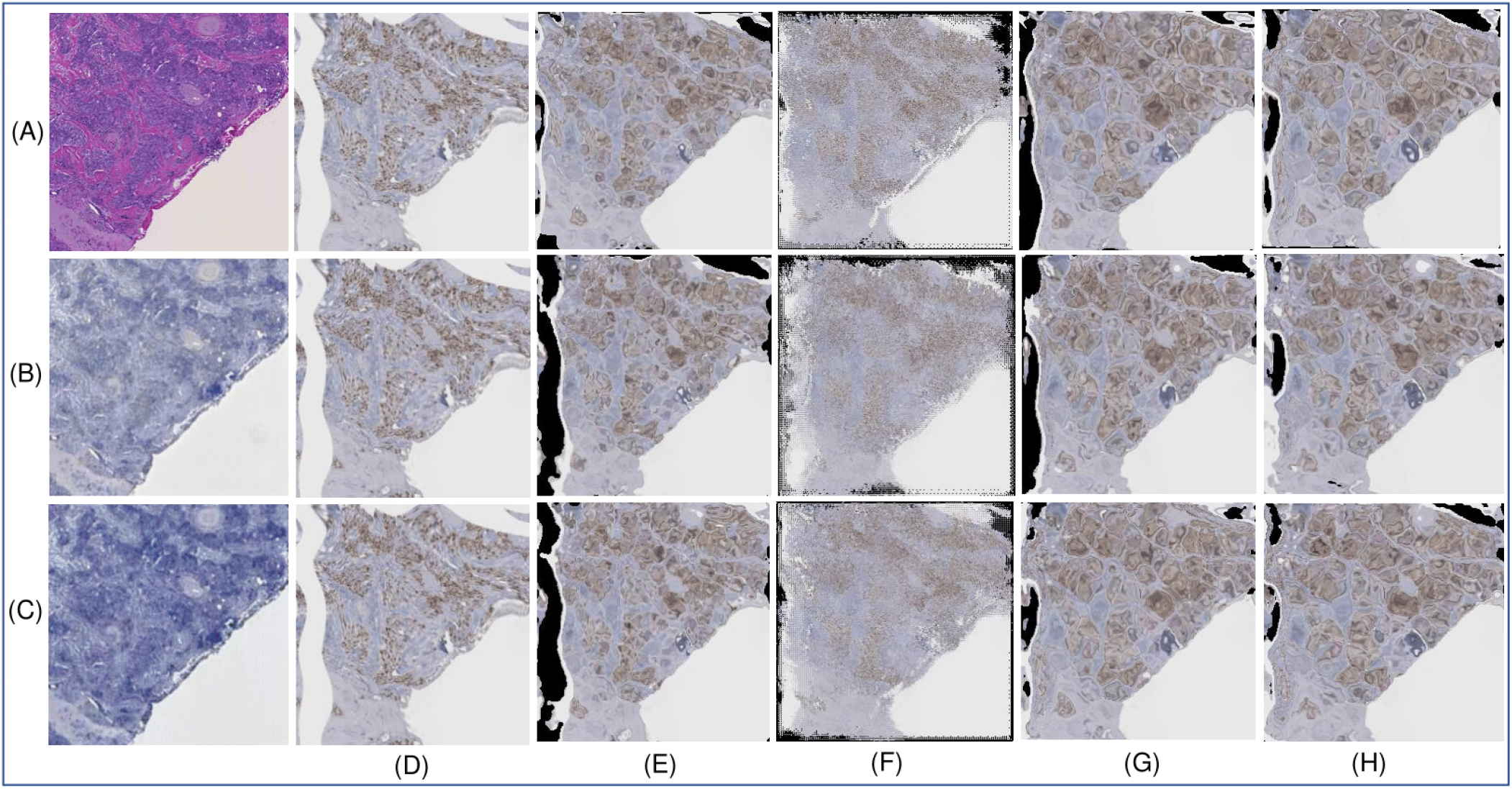
Patch based registration performance with the moving image artificially warped by an elastic deformation. (A)Fixed real H&E image; (B)The resulting synIHC image from (A) by CycleGAN; (C)The resulting synIHC image from (A) by our modified CycleGAN; (D) Moving images after manual transformations; Registration results by (E)DirNet, (F)VoxelMorph Unet, (G)FCN, and (H)our proposed multi-scale FCN.

After method evaluations with manually deformed images, the same methods are applied to ‘real’, ‘syn-1’ and ‘syn-2’ from the testing and validation data, respectively. Representative registration results are demonstrated in Figure 9 and Figure 10, respectively. By visual comparisons, baseline FCN and our proposed multi-scale FCN models perform better than other methods for comparison when images with our modified CycleGAN from the ‘syn-2’ translation result set are used. Additionally, we present quantitative performance evaluation results in Table 2 where Normalized Cross Correlation (NCC), SSIM and Normalized Mutual Information (NMI) are used to report the registration performance. Note our developed multi-scale FCN with the ‘syn-2’ translation result set from the testing data achieves the best performance (bold) by NCC and the second best (underlined) by SSIM and NMI. For the validation data, our developed multi-scale FCN achieves the third best (italics) with ‘syn-2’ by NCC, the second best with ‘syn-2’ by SSIM, and the best value with ‘syn-1’ by NMI. By contrast, the performance of the conventional registration method Elastix is limited due to its over-deformed image outputs (C.F. Supplemental Figure 2). Additionally, all deep learning-based models outperform with ‘syn-2’ than ‘syn-1’ or ‘real’ for the testing dataset, suggesting the efficacy of the enhanced image translation quality by our modified CycleGAN.

**Fig. 9.**
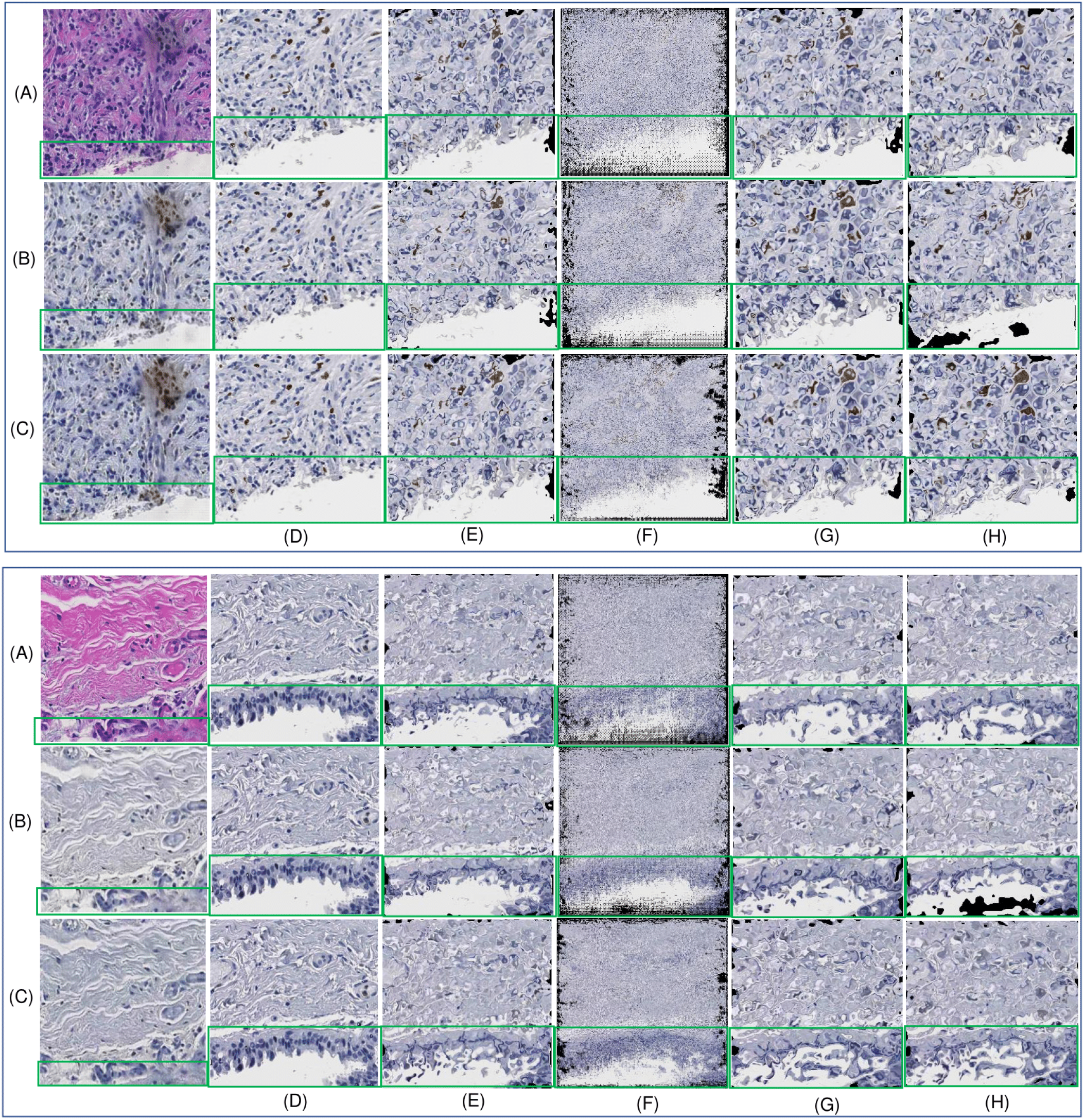
Typical patch registration performance for the testing data. (A)Fixed real H&E image; (B)The resulting synKi67 image from (A) by CycleGAN; (C)The resulting synKi67 image from (A) by our modified CycleGAN; (D)Real Ki67 moving image; Registration results by (E)DirNet, (F)VoxelMorph Unet, (G)FCN, and (H)our proposed multi-scale FCN. Green boxes are used to highlight the registration results.

**Fig. 10.**
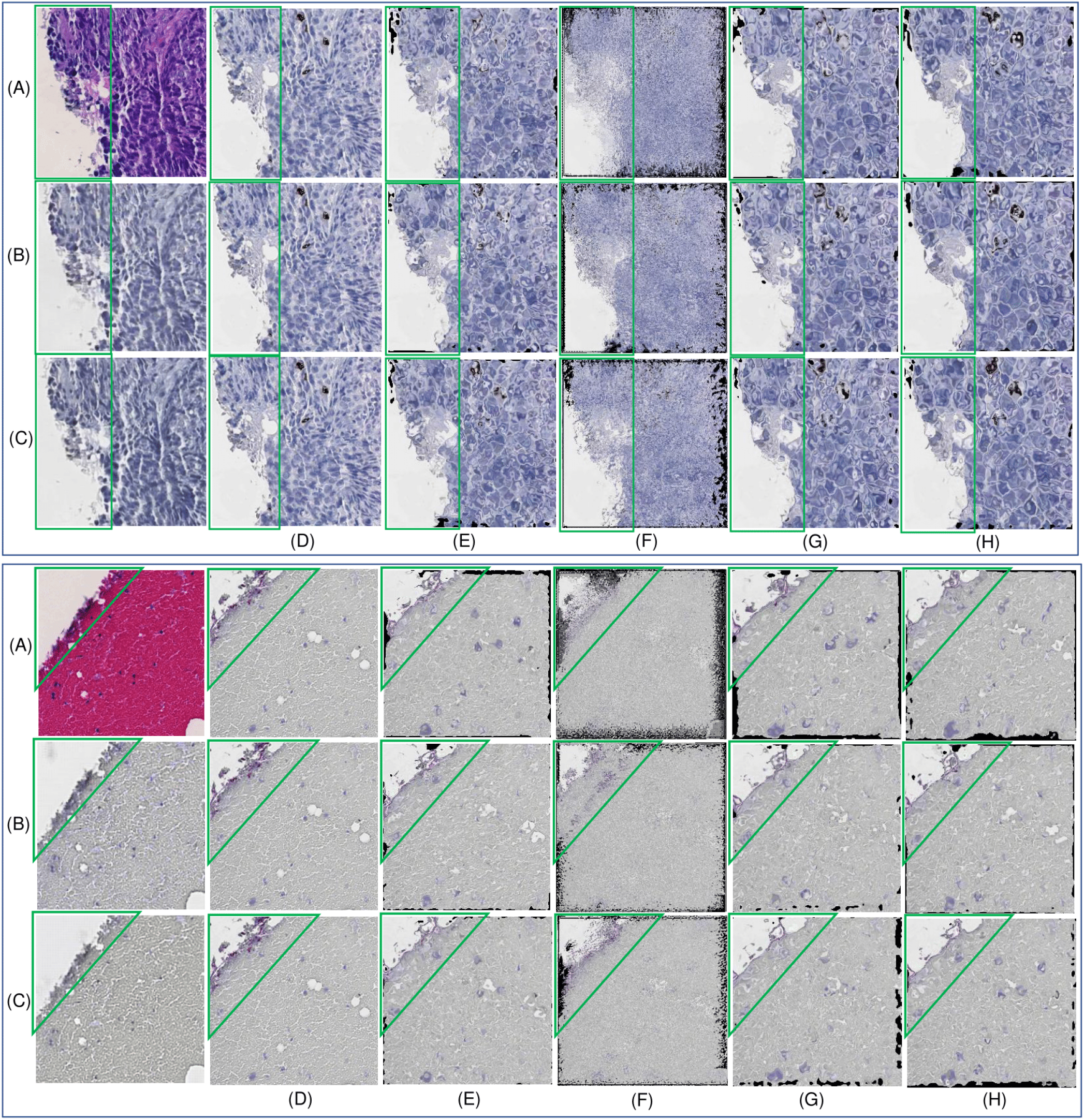
Typical patch registration performance for the validation data. (A)Fixed real H&E image; (B)The resulting synPHH3 image from (A) by CycleGAN; (C)The resulting synPHH3 image from (A) by our modified CycleGAN; (D)Real PHH3 moving image; Registration results by (E)DirNet, (F)VoxelMorph Unet, (G)FCN, and (H)our proposed multi-scale FCN. Green boxes are used to highlight the registration results.

Additionally, two measures of the predicted DVF are used for deformation quality assessment. First, we compute the Jacobian determinant to evaluate the invertibility of the DVF transformation [36]. The Jacobian determinant of the DVF (*V*) at a given point 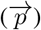 is defined as:

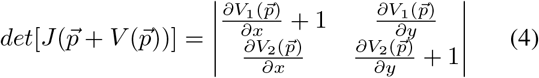

where, 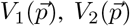 are two components of the DVF 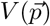 at pixel 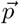. Negative Jacobian determinant at a pixel suggests that the local deformation is not invertible, causing the unrealistic folding artifacts within tissues. A Jacobian determinant either way larger than 1, very close to but above 0, or less than 0 indicates a poor deformation field quality. Suggested by literature, 1 – 3%of negative Jacobian determinant is generally acceptable for a deformable image registration [37]. In Table 3, we present the percentage of negative Jacobian determinant (i.e. folding%), standard deviation, mean, and median of Jacobian determinant for the testing and validation data. Note that percentage of negative Jacobian is zero in all but VoxelMorph Unet method, suggesting the presence of unrealistic deformations by VoxelMorph Unet. This agrees with our visual assessments.

**Table 3.** Percentage of folding, standard deviation, mean and median of the Jacobian of DVFs with testing and validation data for image patch registration.

| Method Name | dataset | Testing Data |  |  |  | Validation Data |  |  |  |
| --- | --- | --- | --- | --- | --- | --- | --- | --- | --- |
|  |  | folding% | std | mean | median | folding% | std | mean | median |
| DirNet | real | 0.0000 | 0.0118 | 0.9999 | 0.9996 | 0.0000 | 0.0131 | 0.9997 | 0.9992 |
|  | syn-1 | 0.0000 | 0.0128 | 0.9997 | 0.9994 | 0.0000 | 0.0121 | 0.9997 | 0.9995 |
|  | syn-2 | 0.0000 | 0.0118 | 0.9997 | 0.9993 | 0.0000 | 0.0124 | 0.9997 | 0.9992 |
| VoxelMorph<br>Unet | real | 1.4054e-04 | 0.1506 | 0.9979 | 0.9971 | 7.6896e-04 | 0.1449 | 1.0024 | 1.0006 |
|  | syn-1 | 1.2716e-05 | 0.1531 | 0.9999 | 0.9884 | 1.2046e-05 | 0.0963 | 1.0025 | 0.9999 |
|  | syn-2 | 2.0077e-06 | 0.1056 | 1.0000 | 0.9947 | 0.0000 | 0.0680 | 0.9973 | 0.9978 |
| FCN | real | 0.0000 | 0.0133 | 0.9999 | 0.9996 | 0.0000 | 0.0156 | 1.0000 | 0.9993 |
|  | syn-1 | 0.0000 | 0.0125 | 0.9997 | 0.9991 | 0.0000 | 0.0146 | 0.9995 | 0.9989 |
|  | syn-2 | 0.0000 | 0.0133 | 0.9997 | 0.9992 | 0.0000 | 0.0144 | 0.9995 | 0.9985 |
| Our Multi-scale FCN | real | 0.0000 | 0.0131 | 1.0000 | 0.9996 | 0.0000 | 0.0141 | 0.9998 | 0.9990 |
|  | syn-1 | 0.0000 | 0.0132 | 0.9998 | 0.9993 | 0.0000 | 0.0126 | 1.0001 | 0.9998 |
|  | syn-2 | 0.0000 | 0.0125 | 0.9996 | 0.9990 | 0.0000 | 0.0128 | 0.9996 | 0.9989 |

The second measure is Curl of the DVF that suggests local circular motions within a local tissue region [37]. A rapidly changing Curl in a DVF suggests an unrealistic deformation. In Figure 11, we overlay the opposite deformation vectors in the associated DVF of a typical image pairs from ‘syn-2’ result set by DirNet, VoxelMorph Unet, FCN and our proposed multi-scale FCN with the corresponding Curl heatmaps. In this Figure, the fixed image in (A) needs to be rotated to match the moving image in (B) denoted by the opposite deformation vectors. Suggested by the deformation vector directions and the Curl magnitudes, the DVF produced by the DirNet model in (C) does not capture such a circulation motion. Additionally, it is noticeable that the DVF from VoxelMorph Unet presents drastically changing vector directions in (D), resulting a high Curl value overall. By contrast, the FCN model (E) and our multi-scale FCN method (F) can produce a smooth DVF as suggested by the quiver plot and the Curl heatmap.

**Fig. 11.**
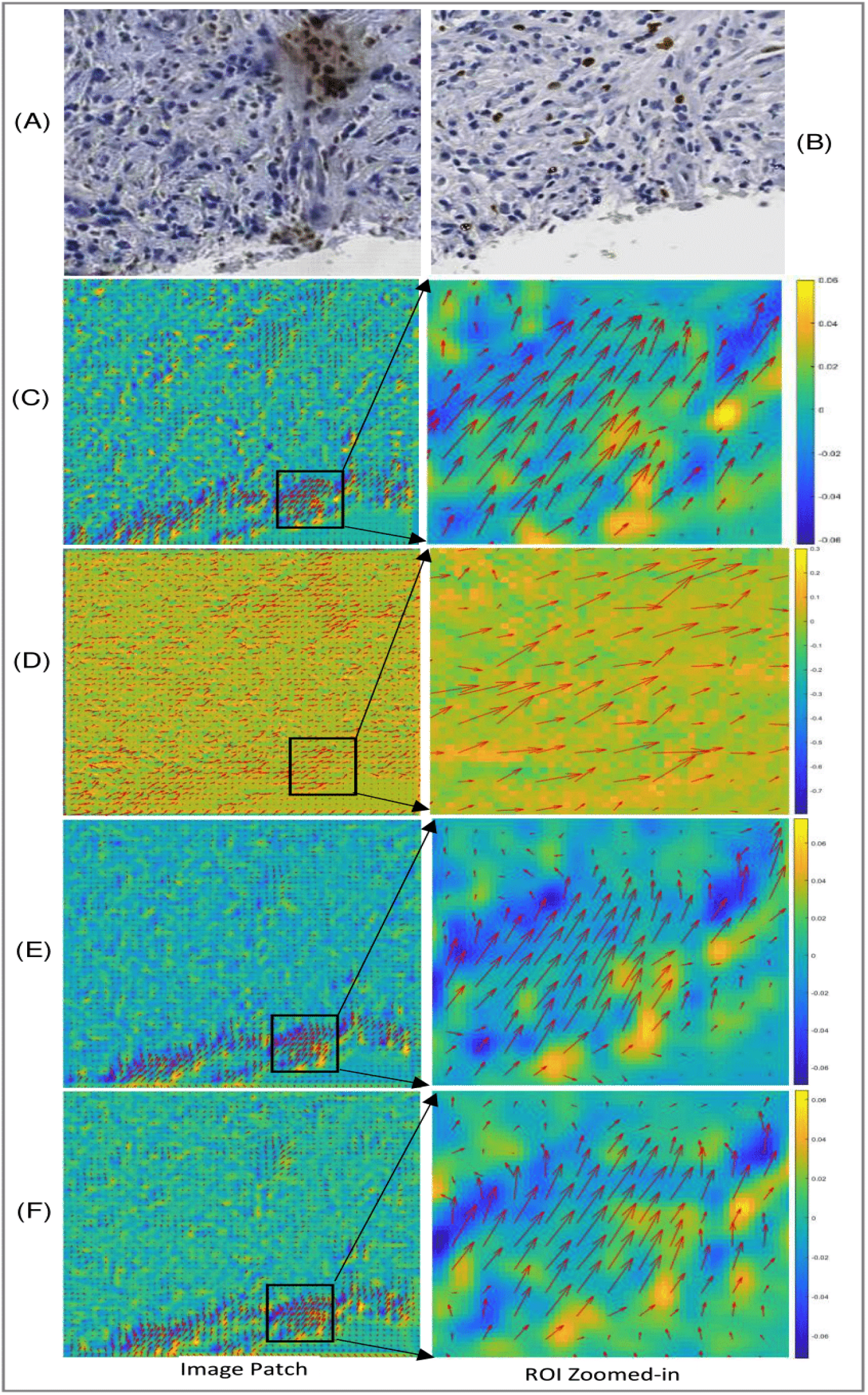
Typical patch-based DVFs overlaid with Curl heatmaps by different methods. (A)Fixed synKi67 image by our modified CycleGAN; (B)Real Ki67 moving image; Quiver plots of DVFs overlaid with Curl values by (C)DirNet, (D)VoxelMorph Unet, (E)FCN, and (F)our multi-scale FCN.

### 3.4. Evaluation of WSI Block Registration

We further evaluate the image registration with WSI blocks. As each WSI block has a size of 8, 000 × 8, 000 pixels, it is partitioned into smaller image patches for the registration process. After individual image patch registrations, they are spatially assembled to generate the registered WSI blocks. In this study, we adopt Dice Similarity Coefficient (DSC) [38], Hausdorff Distance (HD) [39], SSIM, and NCC for WSI block registration accuracy evaluation [40]. For DSC and HD evaluation, eight Region of Interest (ROI) pairs are manually annotated from the testing and validation WSI blocks. The complete evaluation process is presented in Supplemental Figure 3 and the quantitative results are shown in Table 4. Our multi-scale FCN model is compared with both state-of-the-art deep learning based (i.e. DirNet, FCN, and VoxelMorph Unet) and conventional pathology image registration methods (i.e. Elastix, BUnwarpJ [41], Diffeomorphic Demons [42], and WSIReg [6]). Of all methods for comparison, our CycGANRegNet consisting of the modified CycleGAN translation and the multi-scale FCN produces the best performance (bold) by NCC for both testing and validation data. Additionally, our method presents the second best performance (underlined) by DSC for the testing and validation data. By the metric of HD, our method achieves the second best and third best (italics) for testing and validation data, respectively. Although no method produces either the best or second best performance by all metrics for both testing and validation data, our multi-scale FCN model with our modified CycleGAN image translation method presents the best, second best or third best performance in most metrics. Results in Table 4 also suggest that the image translation from H&E to synIHC helps to improve registration performance in most cases.

**Table 4.**
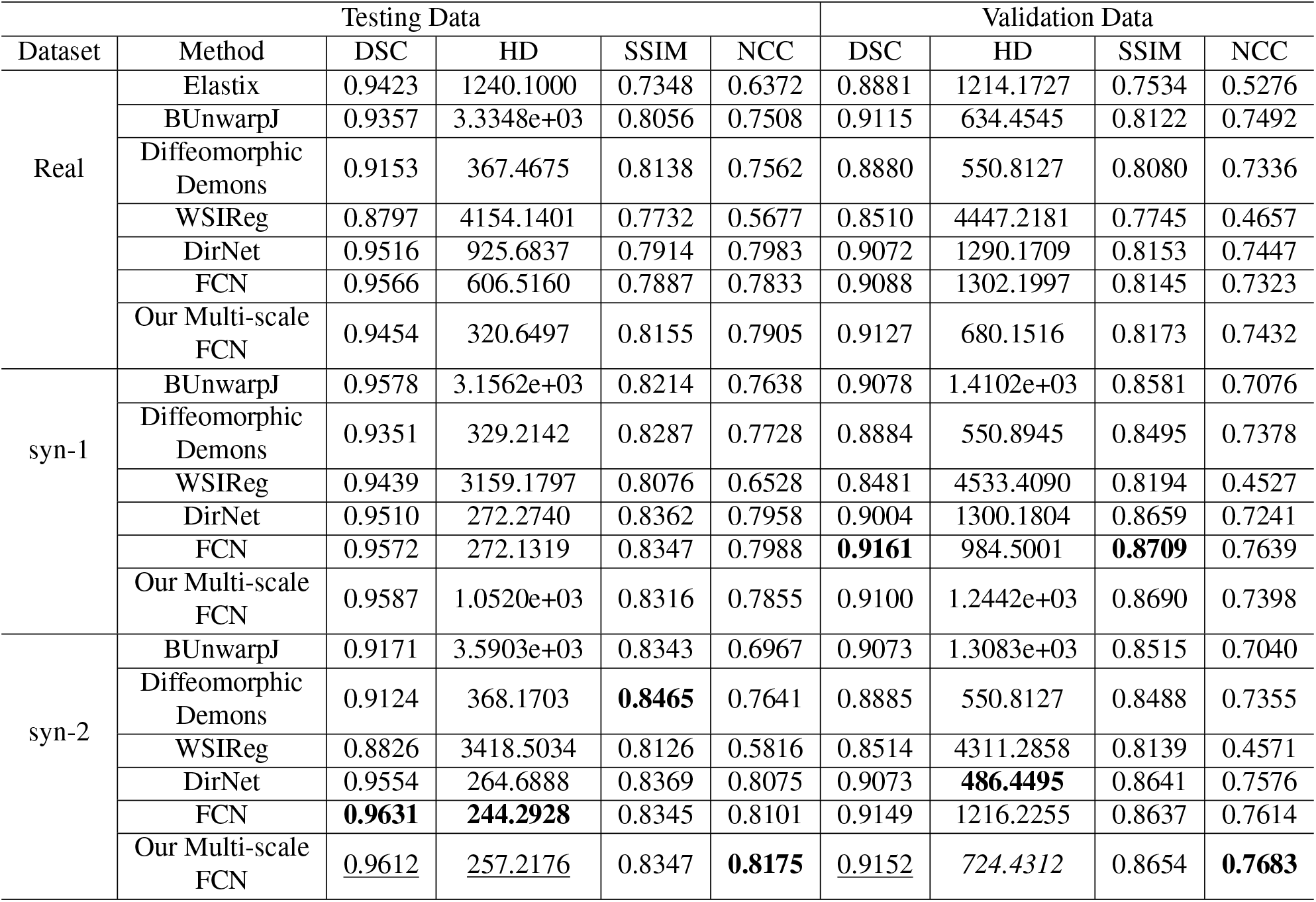
Registration performance of WSI blocks from the testing and validation data.

We present the registration results of WSI blocks from the testing and validation data in Figure 12 and Figure 13, respectively. Additionally, we superimpose a representative WSI block in H&E with opposite deformation vectors in the associated DVF in Figure 14. These vectors suggest the direction and magnitude of deformation at a given location in the H&E block to match the IHC block. Both quantitative and qualitative results suggest that our CycGANRegNet exhibits promising registration performance.

**Fig. 12.**
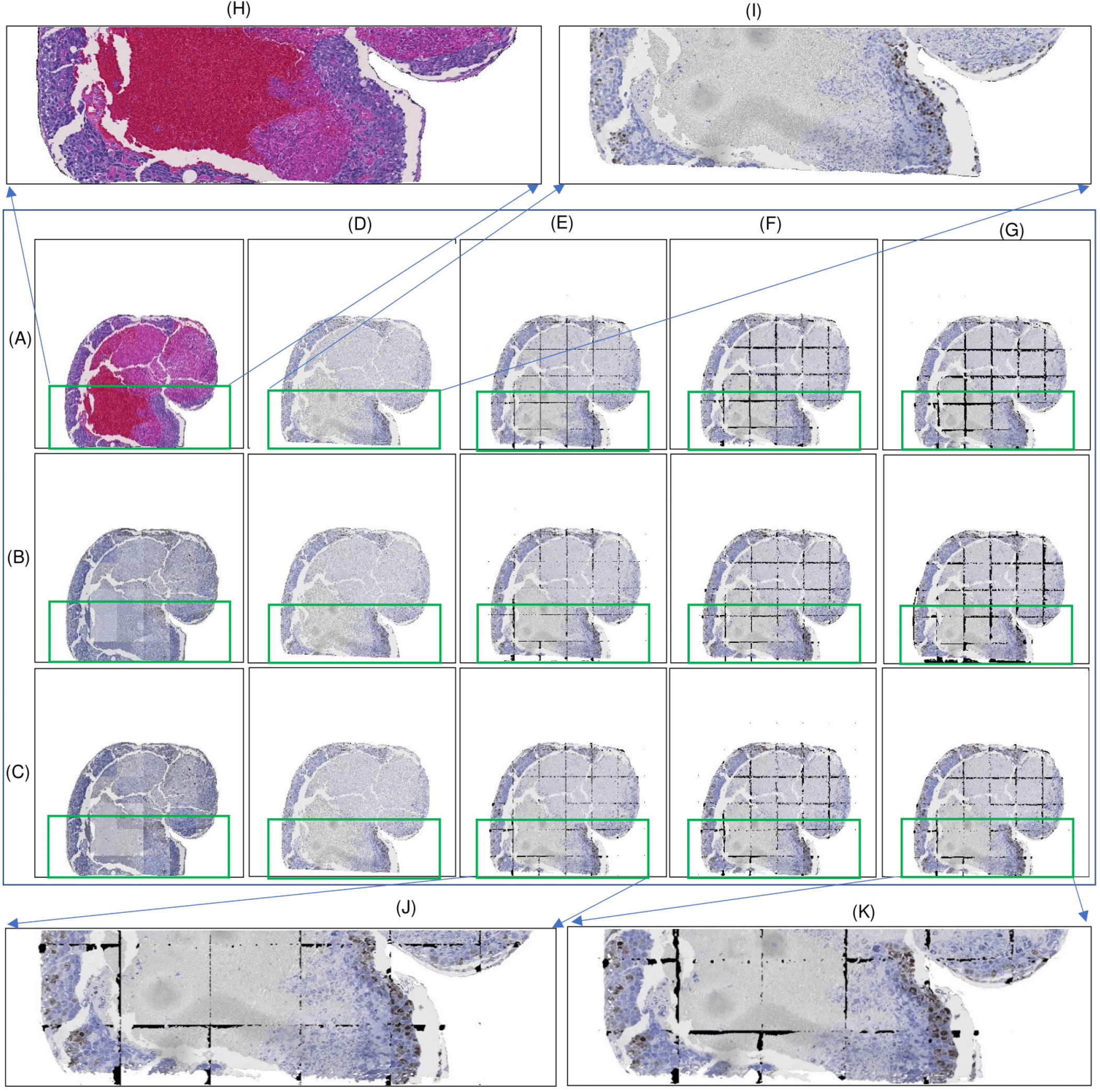
WSI block registration with H&E and Ki67 IHC testing image data. (A)Fixed real H&E image block; (B)The resulting synKi67 image block translated from (A) by CycleGAN; (C)The resulting synKi67 image block translated from (A) by our modified CycleGAN; (D)Real Ki67 moving image block; Registration results by (E)DirNet, (F)FCN, and (G)our multi-scale FCN; (H)Close-up view of a tissue region in (A); (I)Close-up view of a tissue in (D); (J)Close-up view of a tissue in the registered block by DirNet with the ‘syn-2‘ translated dataset; and (K)Close-up view of a tissue in the registered block by CycGANRegNet.

**Fig. 13.**
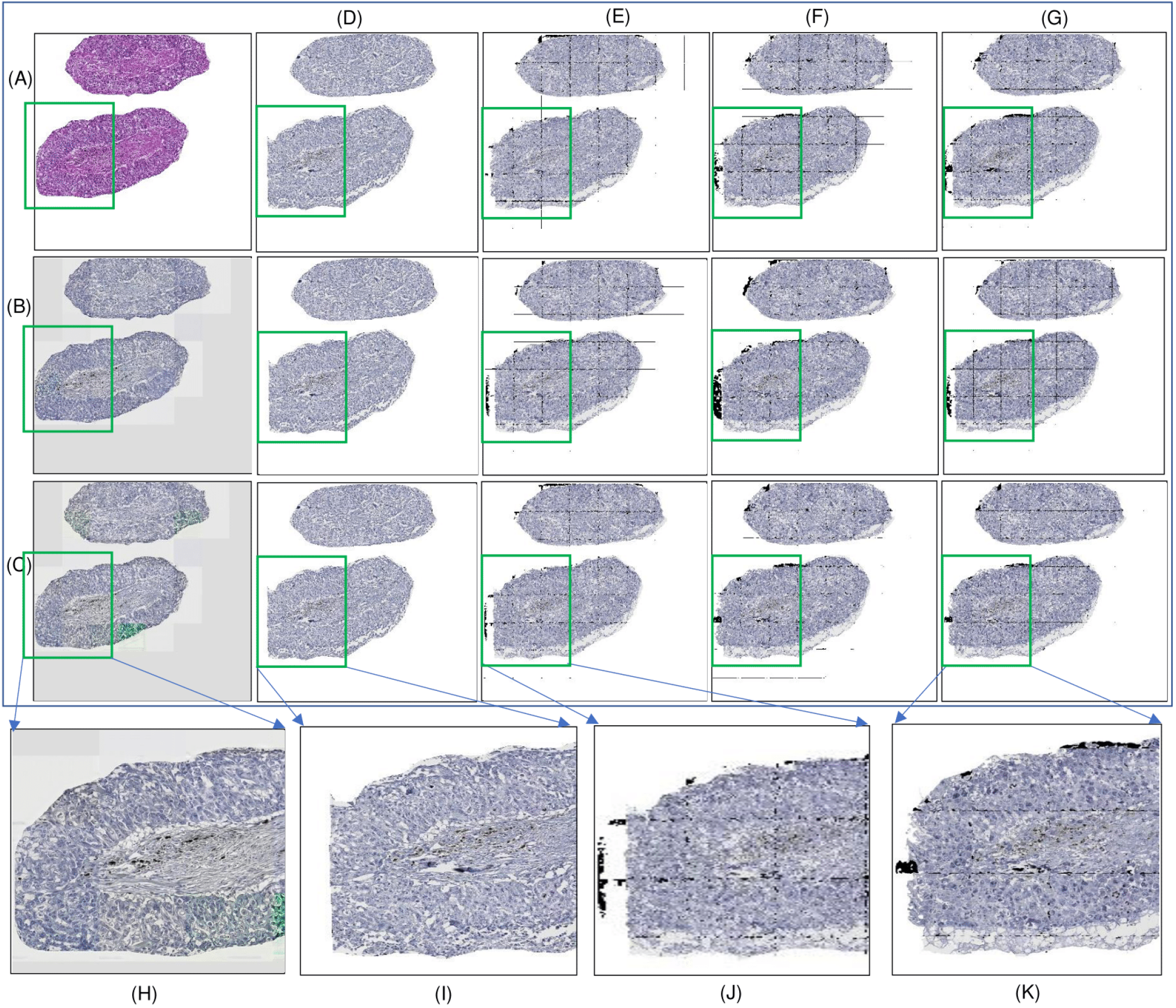
WSI block registration with H&E and PHH3 IHC validation image data. (A)Fixed real H&E image block; (B)The resulting synPHH3 image block translated from (A) by CycleGAN; (C)The resulting synPHH3 image block translated from (A) by our modified CycleGAN; (D)Real PHH3 moving image block; Registration results by (E)DirNet, (F)FCN, and (G)our multi-scale FCN; (H)Close-up view of a tissue region in (A); (I)Close-up view of a tissue in (D); (J)Close-up view of a tissue in the registered block by DirNet with the ‘syn-2‘ translated dataset; and (K)Close-up view of a tissue in the registered block by CycGANRegNet.

**Fig. 14.**
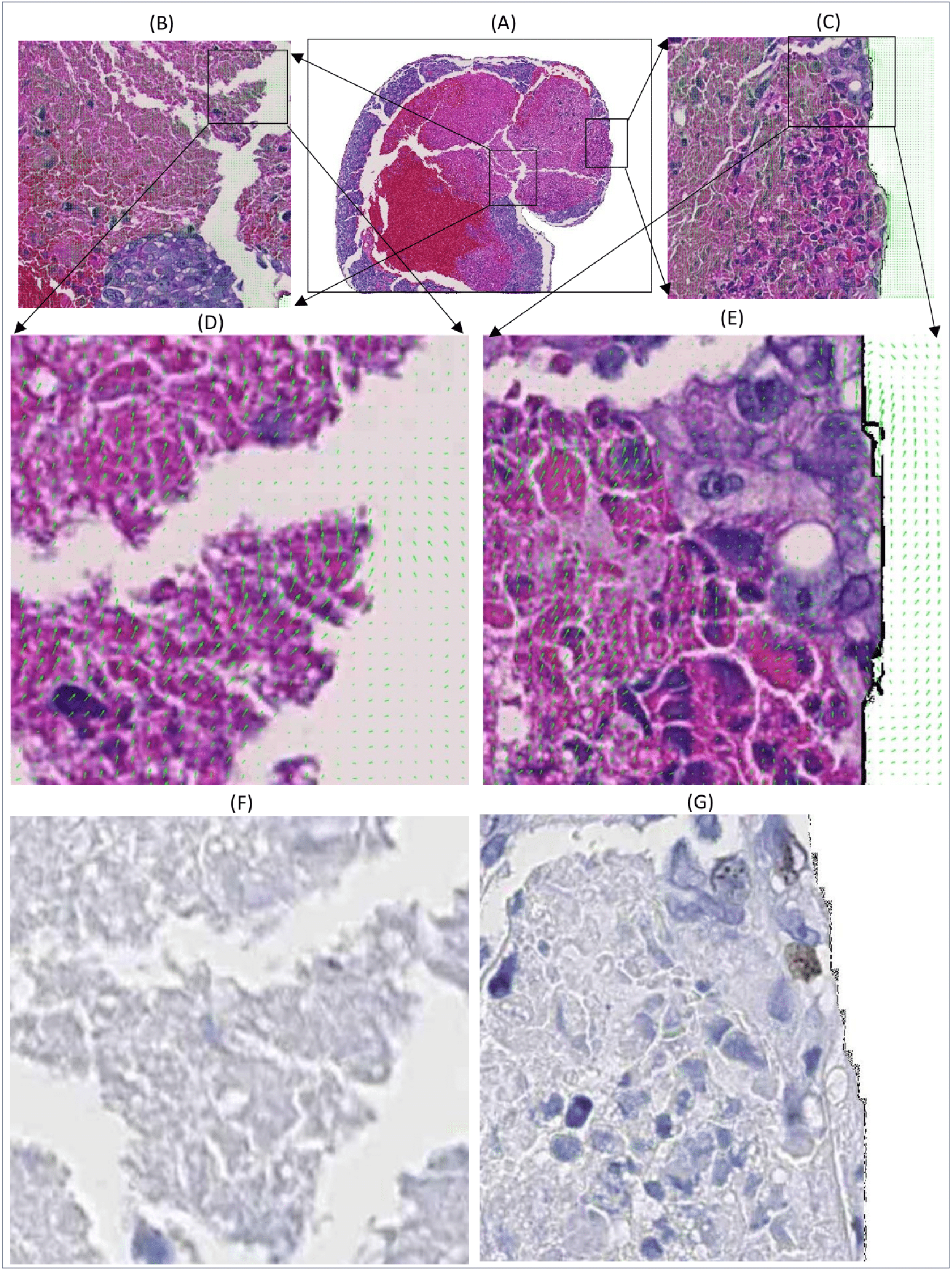
A typical WSI block overlaid with the negative DVF quiver plot. (A)Fixed real H&E block; (B-C) Close-up views of two tissue regions with the negative DVF overlaid; (D-E)Close-up views of (B-C) at a higher resolution; (F-G) Associated regions in the Ki67 IHC moving blocks.

## 4. DISCUSSION

In this study, we develop a two-stage CycGANRegNet model for multi-stained serial WSI registration. By both qualitative and quantitative evaluations, CycGANRegNet demonstrates promising registration performance by multiple metrics. Specifically, the registration component of CycGANRegNet follows a coarse-to-fine multi-scale deformable image registration strategy and optimizes the joint loss at multi-scale levels, leading to a more accurate DVF estimation critical for an enhanced image registration. In Table 2, we notice that performance values (e.g. NCC and SSIM) for the patchbased registration are relatively low as some image patches are extracted from poorly matched WSI block pairs after the pre-alignment step (Supplement Figure S4). For patches from well aligned WSI block pairs after pre-alignment step, such performance values are much improved (c.f. Supplement Figure S5). To test the CycGANRegNet efficacy, we synthetically deform moving image patches by different spatial transformations. In such well controlled experiments, moving images can be well aligned to the fixed images by our method.

Both registration results at the patch and WSI block level suggest the necessity of image translation from one stain to another. Specifically, results in Table 4 suggest that the ‘syn-1’ and ‘syn-2’ translation datasets can produce better registration performance than the ‘real’ dataset in most cases. Additionally, the translated dataset ‘syn-2’ yields better registration performance than the ‘syn-1’ dataset by most metrics, suggesting a better DVF estimation enabled by synIHC images from our modified CycleGAN. Mean-while, our multi-scale FCN with ‘syn-2’ translated dataset consistently produces the best, second best or third best performance values for most metrics, suggesting the superiority of our model to other methods for comparison. In future research, we plan to improve this work by better learning and integrating spatial transformations from the H&E-IHC and synIHC-IHC pipeline for registration. In our current study, we extract non-overlapping patches from each WSI for deep learning model training and evaluation. We will extract overlapping patches to better control the displacement vectors at the patch borders in future.

## 5. CONCLUSION

In this study, we develop a fully unsupervised translation based network CycGANRegNet for H&E and IHC pathology image registration. The resulting synthetic IHC images from the image translation module are aligned to the real IHC counterparts with a multi-resolution approach preserving histology structures at the full image resolution. CycGAN-RegNet can be efficiently trained without ground truth image deformation information. Our method is systematically evaluated and compared with other state-of-the-art methods with testing and validation data. Experimental results at both image patch and WSI block level demonstrate the method efficacy for multi-stained serial WSI registration essential to tissue-based integrative biomedical research studies.

## 6. ACKNOWLEDGMENTS

This research was supported by National Institutes of Health [U01CA242936, R01EY028450, R01CA214928, R01CA239120], National Science Foundation [ACI 1443054, IIS 1350885], CNPq and FAPEMIG agencies.

