## Supplemental file for "Deep Learning Based Registration of Serial Whole-slide Histopathology Images in Different Stains"

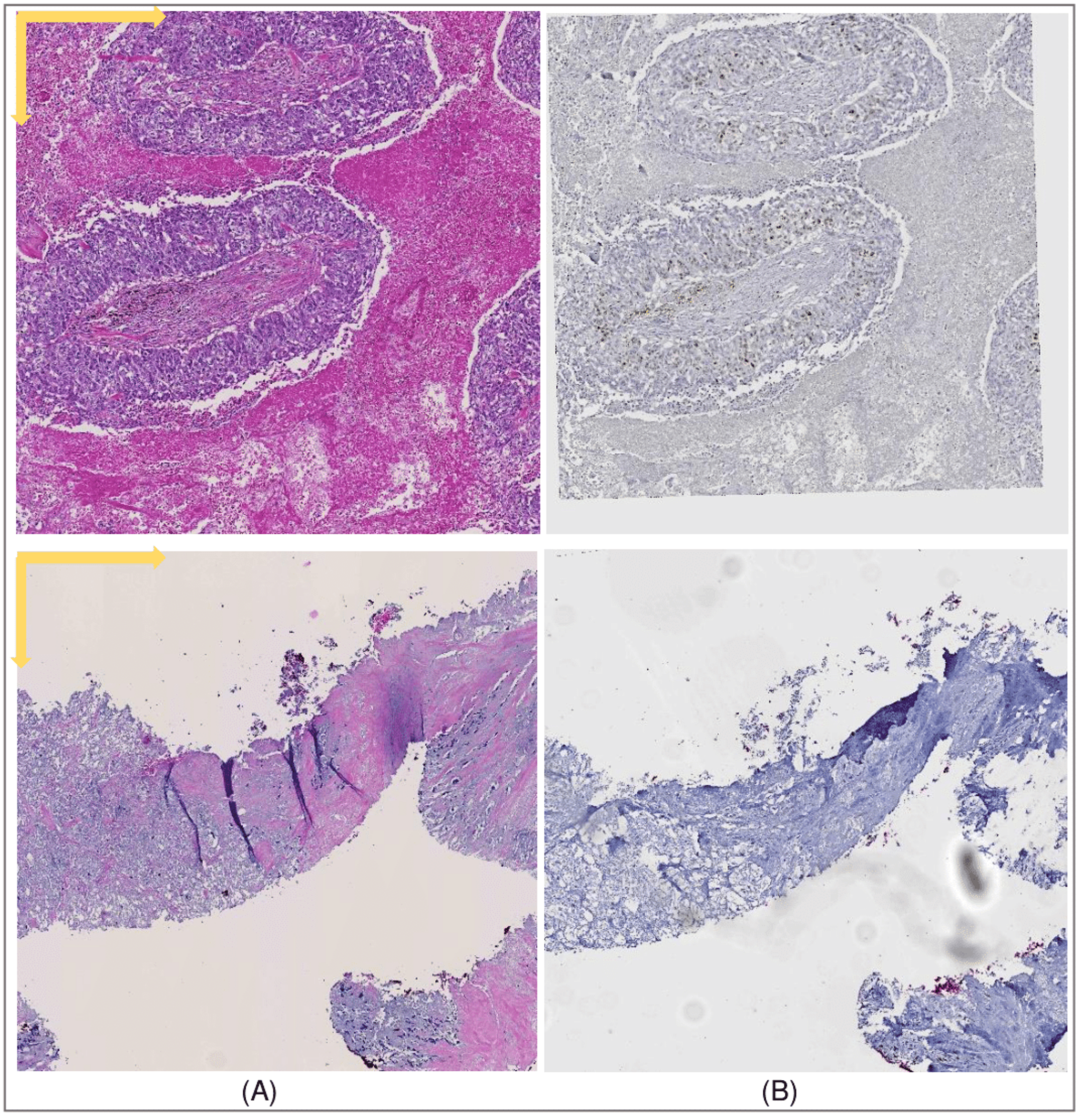


**Fig. 1**. Representative WSI blocks of two serial tissue sections (A) in H&E and (B) of IHC biomarkers.

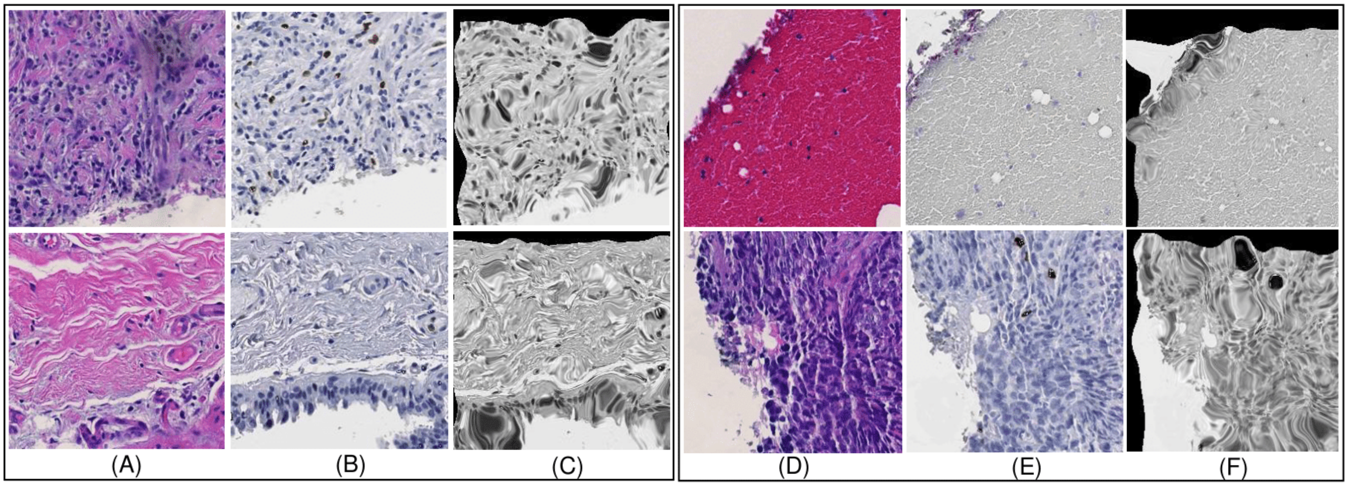


**Fig. 2**. Typical patch registration performance for the testing and validation data by SimpleElastix. (A)Fixed real H&E images; (B)Real Ki-67 moving images; (C)Registration results by SimpleElastix; (D)Fixed real H&E images; (E)Real PHH3 moving images; (F)Registration results by SimpleElastix.

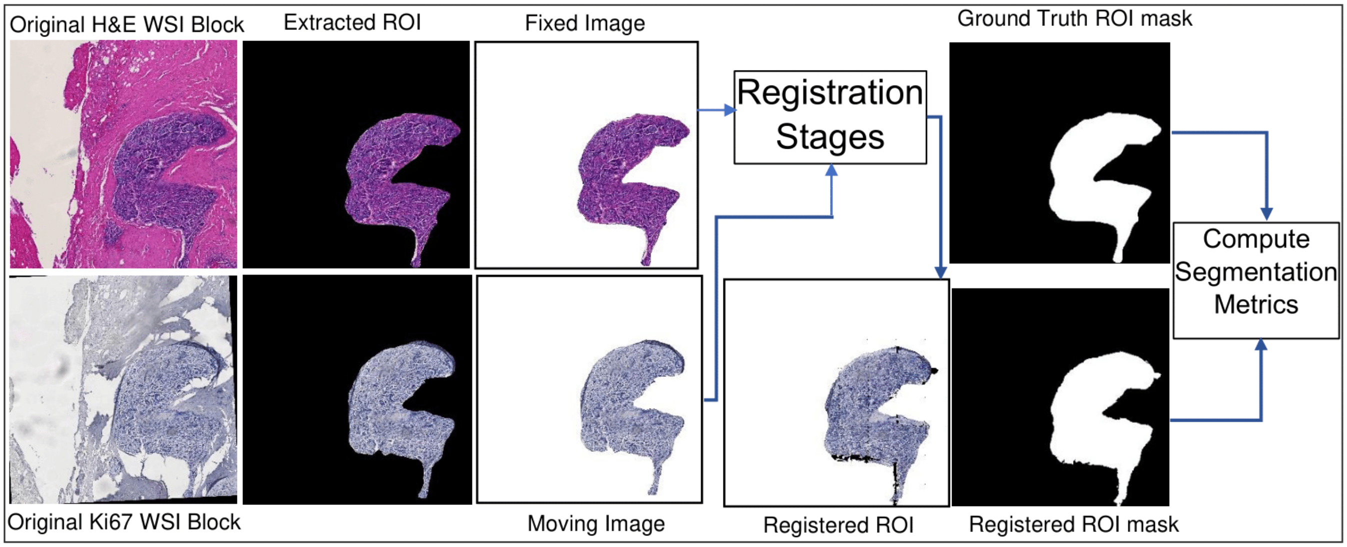


**Fig. 3**. The workflow for DSC and HD computations with ROIs from WSI blocks.

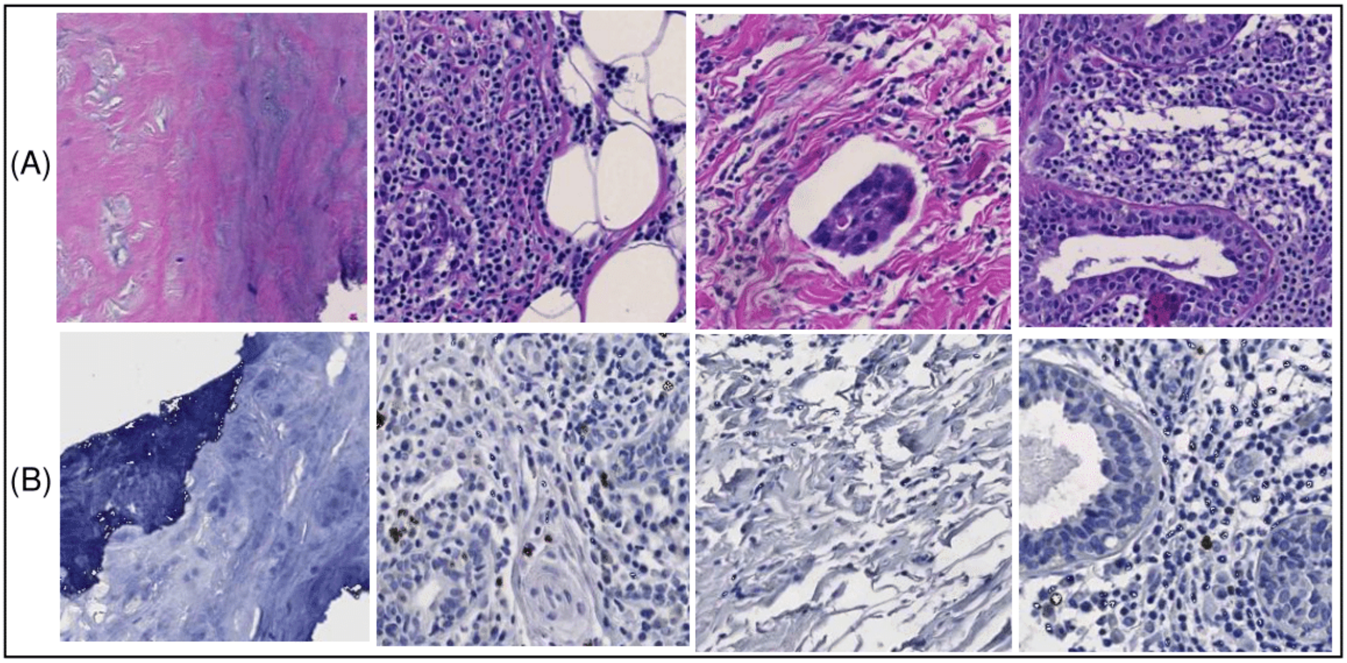


**Fig. 4**. Typical image patch pairs from ill-matched WSI blocks after the pre-alignment step are presented in columns. (A)Fixed H&E and (B)Moving IHC image patches.

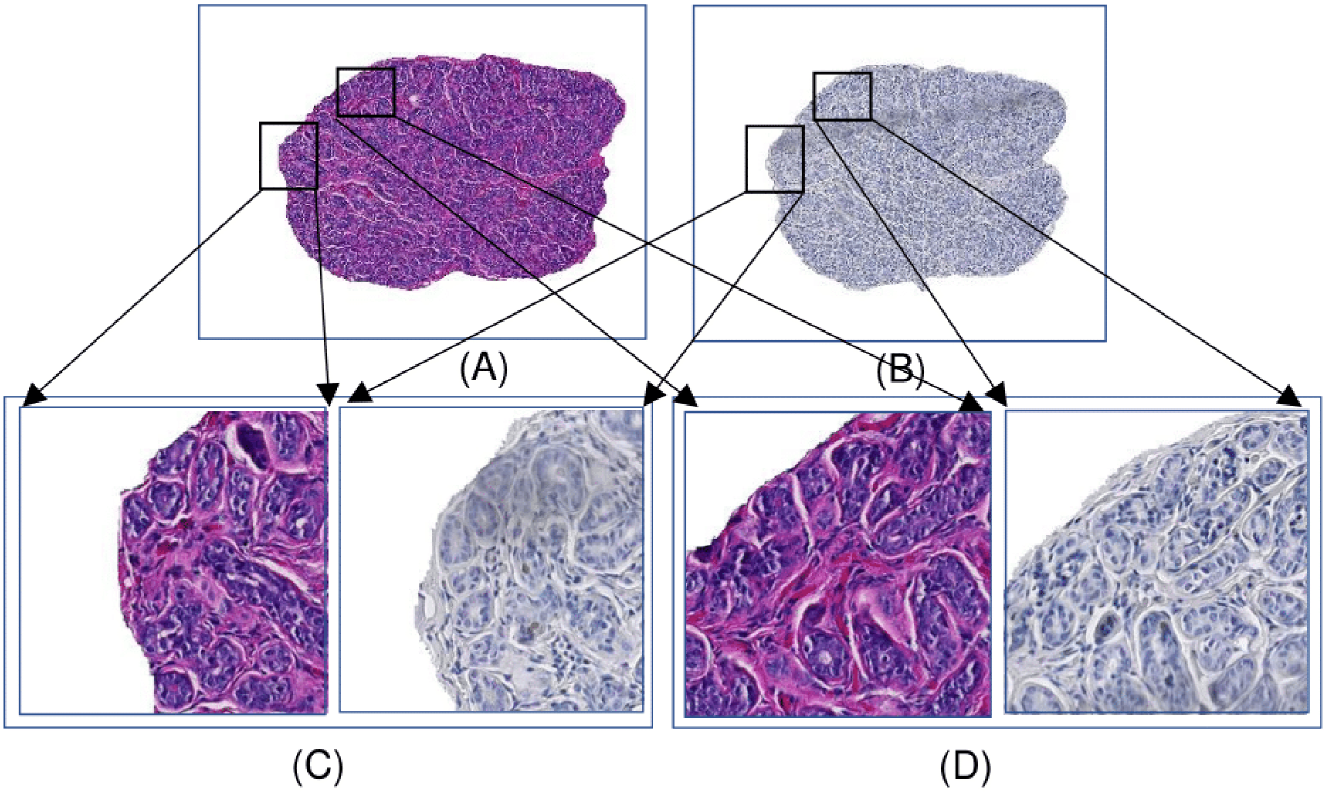


**Fig. 5**. Typical WSI ROI pairs for registration. (A)Fixed H&E WSI ROI; (B)Moving Ki67 WSI ROI; (C-D) Two close-up views of fixed and moving patch-pairs from (A) and (B).
